# Comprehensive identification of fetal cis-regulatory elements in the human genome by single-cell multi-omics analysis

**DOI:** 10.1101/2021.11.02.466852

**Authors:** Hao Yu, Na Ai, Ping Peng, Yuwen Ke, Xuepeng Chen, Yun Li, Ting Zhao, Shan Jiang, Jiang Liu, Lan Jiang

## Abstract

The regulatory programs driving early organogenesis in human is complex and still poorly understood. We performed parallel profiling of gene expression and chromatin accessibility to 28 human fetal tissue samples representing 14 organs in the first trimester. Collectively, we have generated 415,793 single-cell profiles. By integration analysis of transcriptome and chromatin accessibility, we detected 225 distinct cell types and 848,475 candidate accessible cis-regulatory elements (aCREs). By linking regulatory elements to their putative target genes, we identified not only 108,699 enhancers, but also 23,392 silencers elements. We uncovered thousands of genes regulated by both enhancers and silencers in an organ or cell-type-specific manner. Furthermore, our unique approach revealed a substantial proportion of distal DNA elements are transcribed CREs (tCREs), which show both open chromatin signal and transcription initiation activity of non-coding transcript. The landscape of fetal cis-regulatory elements facilitates the interpretation of the genetic variant of complex disease and infer the cell type of origin for cancer. Overall, our data provide a comprehensive map of the fetal cis-regulatory elements at single-cell resolution and a valuable resource for future study of human development and disease.

## INTRODUCTION

Developing and adult human tissues use different cis-regulatory elements but many adult chronic diseases including cancer may have a developmental origin^1–3^. Human fetal development is an exceedingly complex and fascinating process of transforming a single-cell zygote into a fully functioning organism within a mere span of 40 weeks^4^.

And the rudimentary formation of all organ systems raised from three primary germ layers (ectoderm, mesoderm, and endoderm) is completed by gestational week 16^5–7^. A fundamental question is how the precursor cells with the same genetic material differentiate into diverse organs and cell types.

Leveraging single-cell molecular profiling techniques, many efforts have been carried out to explore cell heterogeneity and the development process in one or more organs^8–11^. But the majority of these were focused on transcriptome instead of chromatin states, which may prime to transcription or keep the epigenetic memory to adult cells^12^. Here, we performed massively parallel assays of 5’ single-cell RNA sequencing (scRNA-seq) and single-cell assay for transposase-accessible chromatin and sequencing (scATAC-seq) for 14 human fetal organs. We characterize the chromatin accessibility, transcription initiation activity, interaction of target genes of cis-regulatory elements by integrative analysis of two assays to delineate the regulatory landscape of early organogenesis. Multiple modality rich information did uncover spatiotemporal dynamics of distal DNA elements driving human fetal development and help us further understand epigenomic change underlying disease pathogenesis.

## RESULTS

We collected 1, 2, 13, and 12 fetal organ samples from four human donors ranging from gestational week 8 to gestational week 16 (Fig. 1a and Supplementary Fig. 1a, b). For each sample, we parallelly generated matched 5’ scRNA-seq and scATAC-seq profiles by the droplet-based platform through the optimized protocol. All libraries were prepared with a capture target of 8000 cells.

**Fig. 1.**
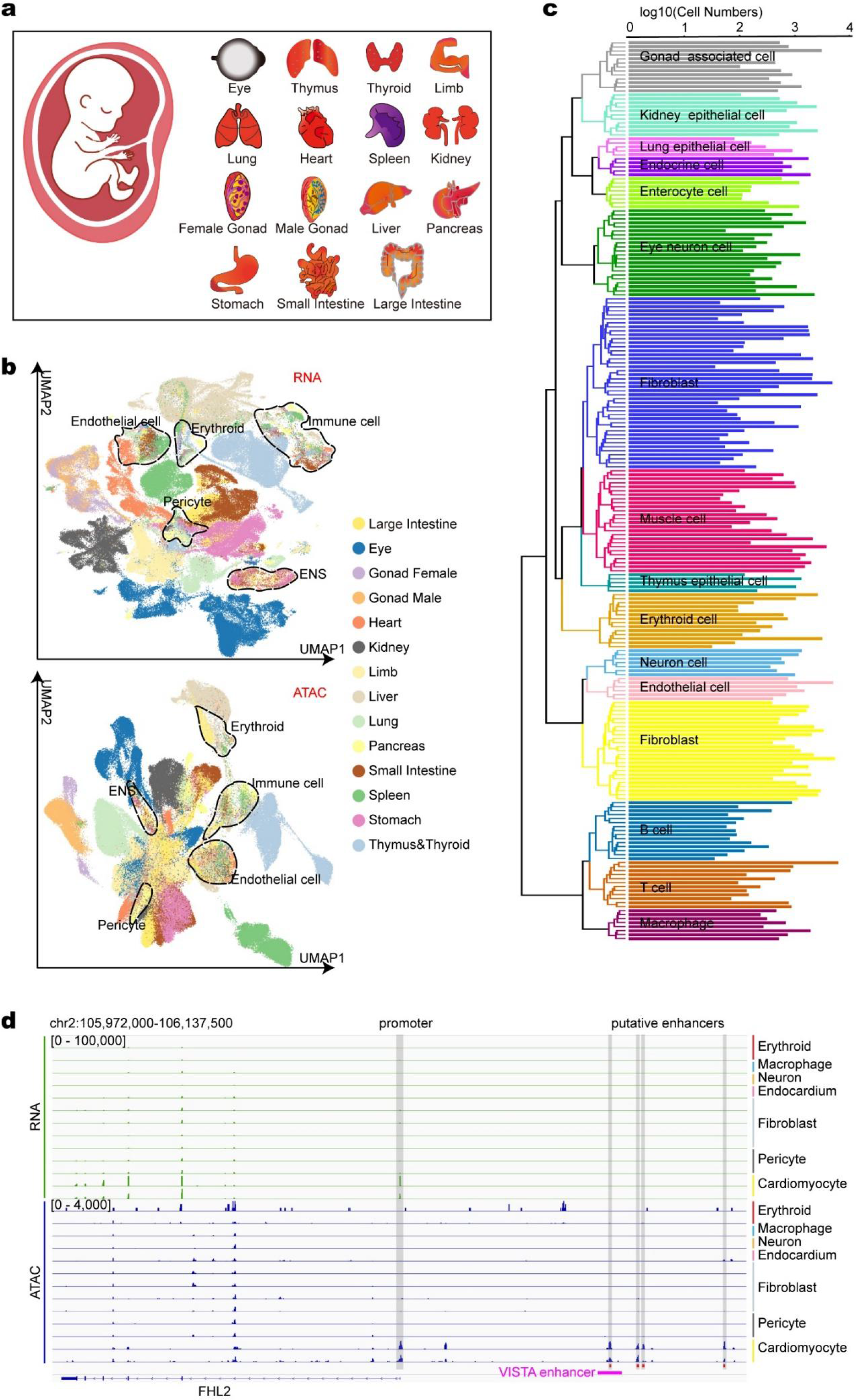
Single-cell transcriptome and chromatin accessibility maps of human early fetus. a, Schematic of collected tissues. b, Upper panel: UMAP embedding of all 185,061 cells from the scRNA-seq data. Lower panel: UMAP embedding of all 212,776 cells from the scATAC-seq data. Each point represents a cell, colored by organ. Some common cell types across organs are outlined. c, Dendrogram showing relationships among 225 cell types. The bar chart on the right represents the number of cells in each cell types in the scATAC-seq data. d, Example locus around FHL2 with differential expression and accessibility across heart-related populations. Shadowed regions highlight the identified cis-regulatory elements.

After quality control, a total of ∼3.1 billion read pairs were retained from scATAC-seq (Supplementary Table 1). These reads constitute 269,920 valid cells. Taken system error into account, we merged multiplet cells about 8% of each library and removed doublet cells about 10% of each library (see Methods). Insert size distribution and TSS enrichment analysis confirms the high quality of our ATAC-seq data (Supplementary Fig. 1c, d). We observed an average level of 9,622 median fragments per cell among 28 samples. Finally, 230,732 high-quality cells with balanced sample sources are used for downstream analysis.

For the matched scRNA-seq for each sample, we applied stringent quality control for the number of detected genes and mitochondrial read counts. Doublets were removed by DoubeltFinder (see Methods). In total, we profiled gene expression in 185,061 individual cells, on average 2,150 genes per cell (Supplementary Fig. 1a and Supplementary Table 1).

### Annotating cell types

Using SeuratV3^13^, we combined single-cell gene expression profiles from all samples and subjected them to batch effect removal and followed by Louvain clustering and UMAP visualization (Fig. 1b and Supplementary Fig. 1e). For the 42 major clusters identified, more than half of them are organ-specific, while others are derived from several organs. C10 (cluster 10) and C34 are mainly from the lung, while C8, C13,

C17, and C24 are a mixture of more than 7 organs. Surprisingly, the mixture clusters represent different common cell types and co-express specific marker genes. For example, C8 expresses endothelial cell markers PLVAP, as well as C13, which expresses enteric nervous system markers ELAVL4 (Supplementary Fig.1 f,g). Because large cell numbers and apparent heterogeneity exist in many of the 42 major clusters, we went into second round Louvain clustering. We identified sub-clusters within each major cluster and got 335 sub-clusters in total. We assign cell type labels to scRNA-seq major clusters and sub-clusters according to known marker genes from literature and HCL references^11^ (Supplementary Table 2). Through 2 rounds of clustering, we were able to identify common cell types across samples while retaining organ-specific cell types.

Next, we transferred cell type labels from 5’ scRNA-seq data to scATAC-seq data within each organ. We computed gene activity scores for scATAC-seq data, aligned cells from scATAC-seq to cells from scRNA-seq in low dimension space, and got a best-fitted label for each cell using ArchR^14^. As some labels have very few cells in scATAC-seq data, we set a cut-off removing transfer results with a low signal-to-noise ratio (Supplementary Fig. 1h) and finally got 225 reliable labels with paired pseudo-bulk profiles of gene expression and chromatin accessibility (Fig. 1c and Supplementary Fig. 1i). To facilitate the exploration of this dataset, we provide an online interface (http://genome.ucsc.edu/cgi-bin/hgTracks?hgsid=1140461557_BMEZ54Vfu607BWs6t5LASYfZT5sj).

### Identify consensus accessible chromatin sites

To construct a map of the cis-regulatory elements marked by chromatin accessibility, we called peaks for each cell type and took the iterative overlap peak merging procedure eliminating redundant peaks using ArchR (see Methods). As a result, the most significant signal in the form of 501bp peaks are caught and a master list of 848,475 consensus accessible chromatin sites are constructed, spanning 14% area of the whole human genome (Supplementary Table 3).

Previous large-scale efforts such as ENCODE3 have mapped open chromatin regions for various tissues/organs and developmental stages mainly based on bulk DNase-seq^15^ or bulk ATAC-seq^16^. However, to what extent, the list of cis-regulatory elements in the human genome is completed is still an open question. We calculated overlaps between peaks we identified and human DHSs of corresponding primary tissues from ENCODE3 (Fig. 2b and Supplementary Fig. 2a). As shown in Venn plots, more than half of the DHSs are detected in our data. And importantly, 153,496 novel peaks are uncovered in our data exclusively. Then, we probed into which tissues/cell types contributing most to the dataset’s specific peaks. The majority of DHSs specific peaks are contributed from adult tissues (Supplementary Fig. 2b) while the majority of scATAC-seq specific peaks are contributed from common cell types such as neurons, macrophages, and endothelial cells, with limited overlap between sub cell types (Fig. 2c and Supplementary Fig. 2c-e). We proposed that common cell types distributing in various organs may be underrepresented in the bulk experiment, while clustering of single-cell data across organs can better capture cis-regulatory elements of those cell types. A significance test in the box plot confirms the above viewpoint (Fig. 2d).

**Fig. 2.**
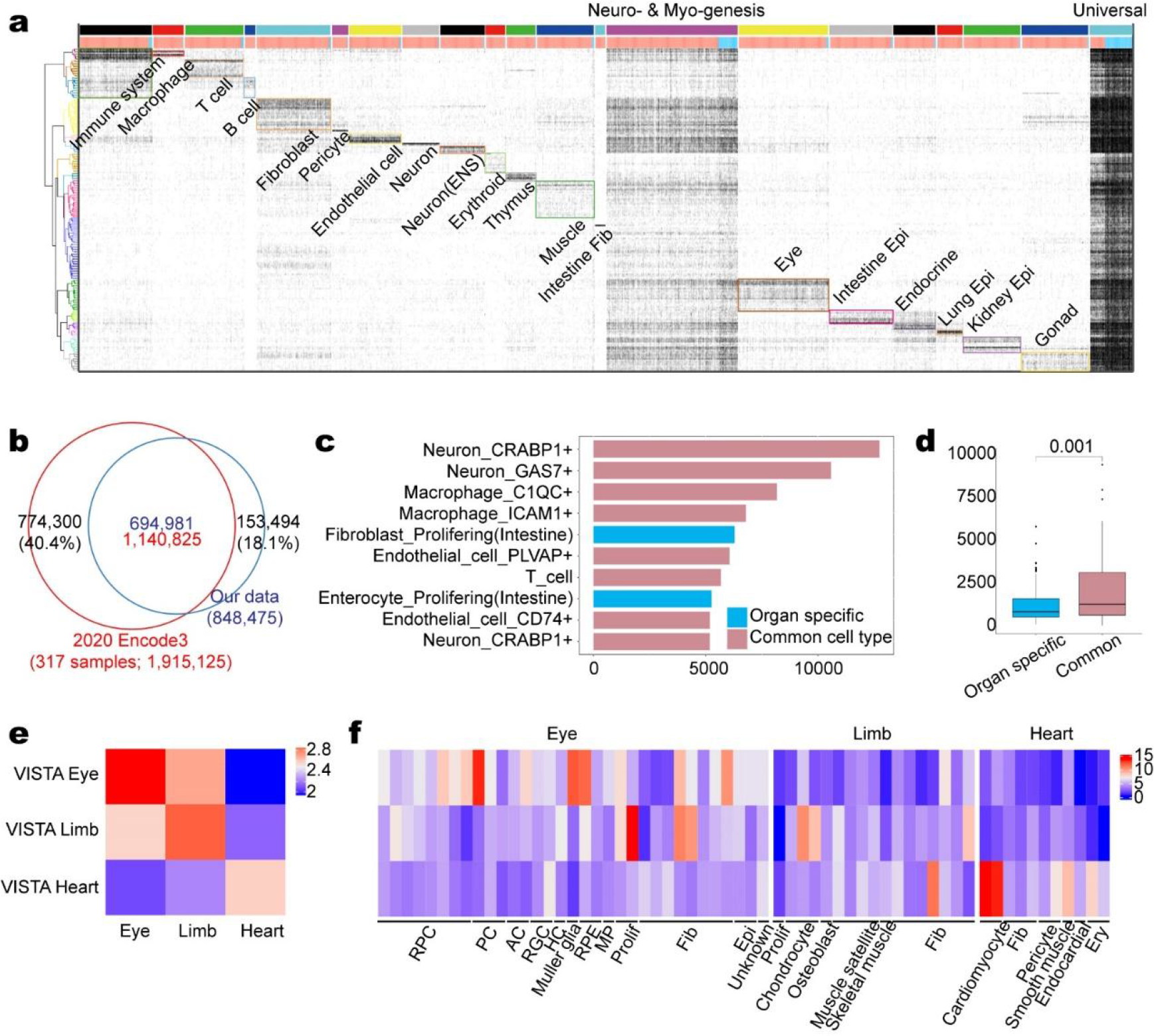
Identifying chromatin accessible sites and patterns in all cell types. a, Chromatin accessibility at 848,475 peaks (x axis) across 225 cell types (y axis). The color code on top represents 21 LSFs. Orangered/deepskyblue color code represnets TSS distal/proximal peaks. b, The overlap between DHSs from ENCODE3 paper and our ATAC peaks. DHSs from corresponding organs/tissues are used for comparison. c, Top 10 cell types that contribute most to ATAC specific peaks (153,494 in Fig 2B). d, Contribution to ATAC specific peaks stratified by two classes of cell types. Boxes denote medians and interquartile ranges (IQRs, 25–75%), whiskers represent 1.5 x IQRs. e, Enrichment for VISTA enhancers within ATAC peaks in the corresponding organ. f, Same as Fig. 2e, but in the cell type level. RPC, retinal progenitor cell; PC, photoreceptor cell; AC, amacrine cell; RGC, retinal ganglion cell; HC, horizontal cell; RPE, retinal pigment epithelium; MP, fetal mesenchymal progenitor cell; Prolif, proliferating cell; Fib, fibroblast; Epi, epithelial cell; Ery, erythroblast.

### Enhancer Validation by Comparison with VISTA database

The VISTA database is a central resource for experimentally validated human and mouse noncoding fragments with gene enhancer activity as assessed in transgenic mice^17^. To know whether the validated enhancers are covered by our results, we drew comparisons on several levels. As showed in the bar plot, over 82% of VISTA enhancers are identified in corresponding organs in our dataset. Besides, about 96% of enhancers are covered without regard to organ sources (Supplementary Fig. 2f). VISTA enhancers are most enriched in the corresponding organ (Fig. 2e), which confirms the tissue specificity of enhancers. More importantly, we can go deep into the cell type level and explore which VISTA enhancers are open in each cell type, expanding our knowledge of enhancers’ function (Supplementary Fig. 2g). For instance, most VISTA enhancers of the heart are open in cardiomyocytes, contributing to the expression of tissue-specific genes like FHL2 (Fig. 1d and Fig. 2f).

### Recognizing the pattern of accessible chromatin regions

To connect accessible chromatin sites with biological cellular contexts, we constructed a binary matrix of 225 cell types × 848,475 peaks in which 1 denotes that the peak is open in the corresponding cell type. After characteristic clustering by rows and columns, we were able to visualize the binary matrix in a fashion of neatly arranged blocks on the diagonal (Fig. 2a). The hierarchical cluster by rows offers the biological information on which cell types are most strongly associated with each peak group. The K-means cluster (K = 21) by columns separates peaks into 21 groups. Based on lineage specificity for each group, we defined peaks in each group as a lineage specifier family (LSF). That is, LSF19 is mainly open in kidney epithelial cells, and may dominant cell differentiation and cell fate decision in the nephrogenesis.

We annotated each LSF with the best-fitted cell lineage based on associated cell types or regulators inferred by motif enrichment. For example, peaks in LSF2 are exclusively accessible among macrophages and are annotated as macrophage-related LSF. Peaks in LSF21 are universally open and over 64% of peaks are proximal to TSS (±1kb), which implies that promoter regions are less dynamic across all cell types. Peaks in LSF13 are open in about half of all cell types and we conjectured its universal function across organs. Motifs most enriched in LSF13 include Atoch1/Tcf12/NeuroG2 and Tcf21/MyoD/Twist2, all of which are helix-loop-helix (HLH) transcription factors and act as key regulators of neurogenesis, myogenesis, and osteogenesis^18^.

### Developmental dynamics of chromatin accessibility

To decipher molecular regulation mechanisms underlying LSFs, we sought to explore chromatin accessibility dynamics within each LSF and transcription regulators in lineage differentiation. Taken LSF19 as an example, we adopted an iterative strategy taking cell types of kidney epithelial and repeating K-means clustering (K = 10) to identify sub pattern of accessible chromatin states (Fig. 3a). And we denote subcluster 3 of LSF19 as LSF19.3. This process produced informative substructures and uncovered a huge difference between progenitor cells (cap mesenchyme, CM) and differentiated cells (primitive vesicle, PV; proximal tubules, PT; Loop of Henle, LoH; distal tubules, DT).

**Fig.3.**
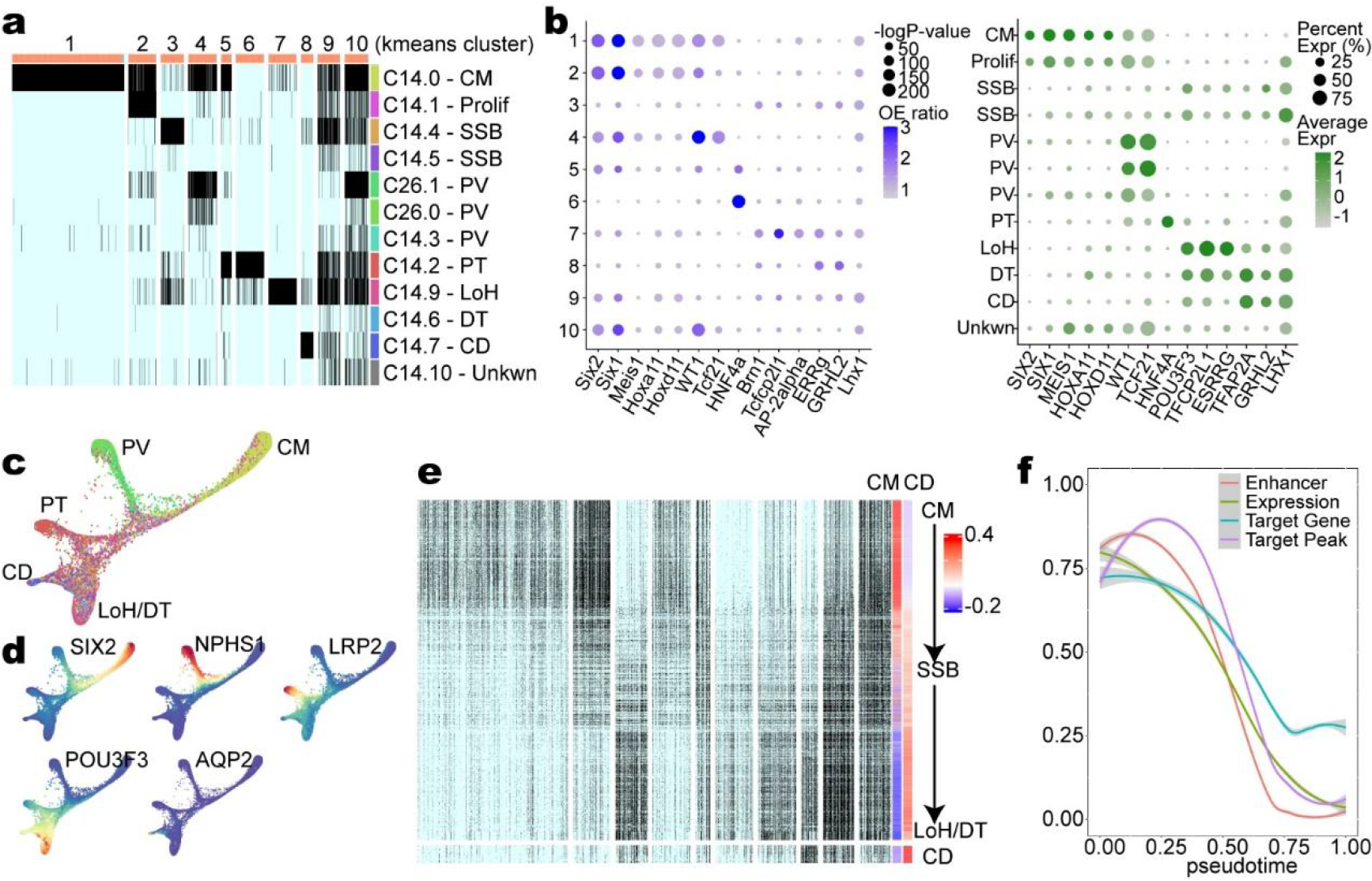
Dynamics of open chromatin and driving transcription factors in nephrogenesis. a, Sub-patterns of chromatin accessible states in G19 from Fig. 2a. All cell types are kidney epithelial cells. CM, cap mesenchyme; Prolif, proliferating cells; SSB, S-shaped body; PV, primitive vesicle; PT, proximal tubules; LoH, Loop of Henle; DT, distal tubules; CD, collecting duct. b, Left panel: Motif enrichment among 10 K-means clusters. Right panel: Expression level of transcription factors among different cell types. The motifs and transcription factors are corresponding in position. c, UMAP embedding of all 12,652 cells from the scATAC-seq data, colored by cell type in Fig. 3a. d, Normalized gene activity score level of 5 marker genes. e, Continuous change of chromatin accessibility states along differentiation of loop of Helen/distal tubule. Each row represents a cell, which is ordered by pseudo-time. The bottom part is from collect ducts as a reference. f, Dynamics of SIX2 expression, chromatin accessiblility of its upstream and downstream peaks and downstream gene expression.

The progenitor cells (CM) seem to have the most open chromatin states. Along with lineage differentiation, a lot of sites like LSF19.1/2 are turned off and other function-relevant regions like LSF19.6/7 are opened while some open states like LSF19.9/10 are maintained. The chromatin accessibility states are modified in a branch-determined way.

Furthermore, we found that motif enrichment was consistent with corresponding TF expression in each cell type (Fig. 3b). LSF19.1/2 were enriched with Six2/Six1 motifs and high expression of SIX2/SIX1 also appeared in progenitor cells like cap mesenchyme. This suggests that transcription factors are responsible for establishing and maintaining open chromatin states.

Next, we performed trajectory inference analysis to resolve lineage differentiation at the single-cell level using scATAC-seq data (Fig. 3c, d and Supplementary Fig. 3). We were surprised to find that chromatin accessibility of LSF19 (only 46838 peaks) has sufficient information to distinguish different cell types and underlies differentiation order, which means PV emerges before other parts in the timeline. The DT and collecting duct (CD) are the final two segments of the kidney nephron with the function of ions absorption and water reabsorption. However, the distal cells of the comma-shaped body (precursor of DT) invade the proximal tip of the UB (progenitor of CD) and fuse to form one continuous P/D axis at early stages. We captured a continuous reprogramming process along with the differentiation to DT at the single-cell level (Fig. 3e). The converge suggests that spatial organization or local function may be a more deterministic factor in chromatin accessibility states compared to cell origin. To understand how transcription factors, help to maintain cell states and play a role in lineage differentiation, we made an in-depth investigation on SIX2, which maintains cap mesenchyme in an undifferentiated state^19^. We found SIX2 as a transcription factor can also target the putative enhancer of SIX2 itself to positively regulate SIX2 expression. Then, we inferred the target genes of TF based on the association of TF target peaks. We found that the dynamics of chromatin accessibility of target peaks and expression of target genes of SIX2 have the same trend as SIX2 expression (Fig. 3f). This suggests that the dynamic of open chromatin states is driven by the expression and function of transcript factors, while cis-regulatory elements regulate gene expression in a forward way.

### Linking regulatory elements to cognate genes

We next asked how distal regulatory elements regulate gene expression. Peak co-accessibility is often used to predict enhancer-promoter interactions^20^. However, the accessibility of ubiquitous opened promoters is usually moderately correlated with gene expression. Therefore, we leveraged the gene expression data and created a correlation-based map between chromatin accessibility peaks and their cognate genes directly (see Methods).

Using correlation analysis, we identified 155,620 positive peak-to-gene links (associated with 108,699 peaks and 12,783 genes) and 34,287 negative peak-to-gene links (associated with 23,392 peaks and 7,628 genes) (Supplementary Table 4). Then we defined positive links as putative enhancer-gene pairs and negative links as putative silencer-gene pairs. For example, FHL2 plays an important role in cardiomyocyte differentiation by negatively regulating the calcineurin/NFAT signaling pathway. And we found the putative enhancers of FHL2 are exclusively open in two sub cell types of cardiomyocytes, confirming the accuracy of our results (Fig. 1d).

### Comparison with ReSE-identified silencers

Pang and Snyder devised a lentiviral screening approach^21^, the repressive ability of silencer elements (ReSE), to systematically identify silencer regions in human cells. They assayed on K562, PMA-treated K562, and HepG2 cell lines, and identified a total of 5472 non-overlapping silencers. To validate our data, we compared our correlation-based silencers and ReSE-identified silencers and found an overlap of 174 silencers. chr5:171602285-171602785 and chr19:48763298-48763798 are two examples with different distributions in 225 cell types (Fig. 4a-c). The former shows a sharp decline in expression when the accessibility of the silencer reaches a level of 0.2, and the latter is much milder with a downward tendency. Based on the sharp decline or not, we can classify silencers into strong silencers or weak silencers (see Methods). These two classes may underline two mechanisms: a switch way through repressed epigenetic states to turn on or off target genes (strong silencers), and a competitive way through transcriptional machinery interactions (weak silencers).

**Fig. 4.**
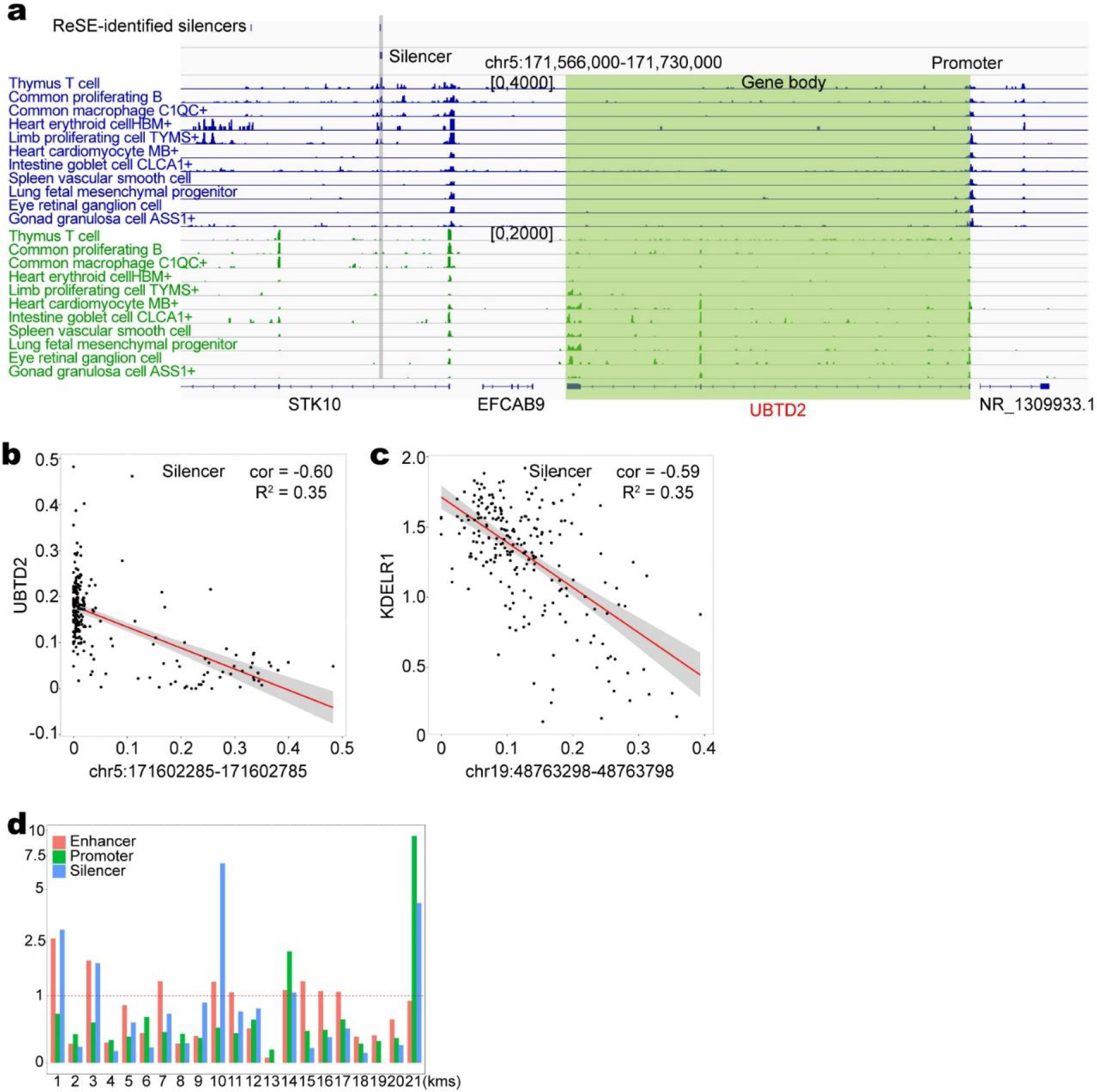
Comparison with ReSE-identified siencers. a, Example locus around UBTD2 with annotated cis-regulatory elements on the top. Cell types are ordered according to the accessibility level of the silencer identified in both study. b, Scatter plot demonstrates the silencer’s accessibility level (x axis), along with UBTD2 expression level (y axis) of each cell type, related to Fig. 4a. c, Scatter plot demonstrates the accessibility level of another ovelaped silencer (x axis), along with UBTD2 expression level (y axis) of each cell type. d, Enrichment for all annotated cis-regulatory elements in different peak groups from Fig. 2a.

To take advantage of our large-scale data, we further predicted target genes for ReSE-identified silencers. 2,113 silencers have at least one neighboring negative correlated gene. Our data and analysis can add complementary information to experimentally verified silencers in whole organism scales (Supplementary Table 5).

### Adversarial regulation on the same gene

To investigate the relationship between our classification of cis-elements and 21 LSFs, we calculated enrichment for each category of cis-elements (Fig. 4d). Interestingly, LSF1, LSF3, LSF10 are enriched with both silencer and enhancers, and they are all related to the hemopoietic system, which underscores a complicated regulatory fashion during hematogenesis, which is consistent with recent report^22^.

Although correlation analysis is based on one peak to one gene, the real situation is that multiple cis-elements cooperatively or competitively regulate the same gene in a cell-type-specific manner. We found a total of 6,091 genes which are the targets of both putative enhancers and silencers (Supplementary Table 6) and focused on a set of 94 genes identified at the whole organism level. Integrated genes expression and open chromatin information allow us better resolve the complexity of regulation (Fig. 5a-c).

**Fig. 5.**
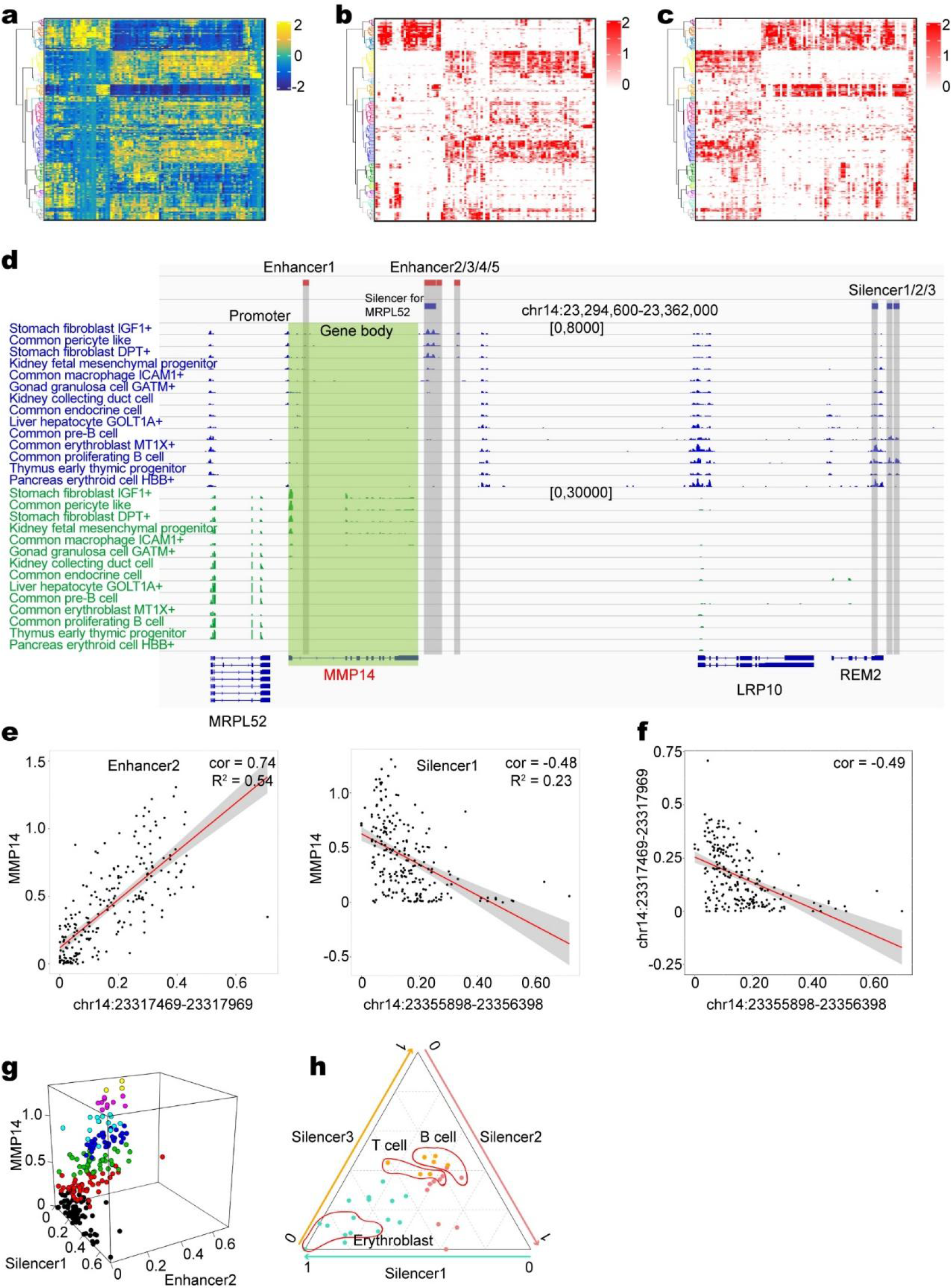
Combinational regulation of positive and negative cis-regulatory elements on the same gene. a-c, Heatmaps showing the combinational regulation on the same gene. (a) Relative gene expression levels. Each column represents a gene. Each row represents a cell type. (b) Relative chromatin accessibility levels of enhancers. (c) Relative chromatin accessibility levels of silencers. Each column represents a peak associated with corresponding gene in (a). Median value is used to represent multiple peaks linked to the same gene. d, Example locus around MMP14 with annotated cis-regulatory elements on the top. Cell types are ordered according to expression level of MMP14. e, Scatter plot demonstrates the peak accessibility level (x axis), along with MMP14 expression level (y axis) of each cell type. Left is enhancer 2 and right is silencer 1 from Fig. 5d. f, Scatter plot demonstrates the accessibility level of silencer 1 (x axis), along with the accessibility level of enhancer 2 (y axis) of each cell type. g, 3D scatter plot showing the relationship among the accessibility level of enhancer 2, silencer 1 and gene expression of MMP14. h, Ternary plot showing the silencer preference among different cell types. Only cell types with normalized expression level less than 0.20 are plotted.

Of the 161 silencers, the majority are open in the hemopoietic system, which is consistent with the cis-regulatory elements & peak group enrichment analysis (Fig. 4d and Supplementary Fig. 4a, b). In line with expectations, the pattern of accessibility of the enhancer is almost the same as the gene expression, while the pattern of accessibility of the silencer is the opposite (Fig. 5a-c). The accessibility pattern of enhancer and silencer of the same gene are mutually exclusive and have a negative correlation. The underlying mechanism will require further investigation.

We next made an in-depth study on one gene, MMP14 (Fig. 5d), whose encoded protein are involved in the breakdown of extracellular matrix in normal physiological processes, such as embryonic development, reproduction, and tissue remodeling, as well as in disease processes, such as arthritis and metastasis. In our dataset, fibroblasts from different organs have high level expression while erythroid cells and immune cells have low level expression. In the track plot, silencers are from close to open and enhancers are from open to close with the decrease of expression level (Fig. 5d). There is a cliff-like change when the accessibility level of silencer 1 reaches the critical point of 0.3, which suggests a switch of regulatory modules (Fig. 5e). When under the critical point, the accessibility of enhancer 2, as well as the expression of MMP14, is highly variable, and enhancer 2 determines the expression level (Fig. 5e-g and Supplementary File 10). Once reaching the critical point, both enhancer and gene transcription is silenced. Mutually exclusiveness of chromatin accessibility between enhancers and silencers uncovers two regulatory modules, functioning in part of cells antagonistically. We further probed into the silencer preference among different cell types. The ternary plot indicates that silencer 1 functions alone in erythroblast, while silencer 2/3 are co-accessible and functional in B cells and T cells (Fig. 5h). The cis-element selection may emerge along with the cell fate decision.

While the silencers in the above example are all strong silencers, we get quite curious about what if one gene is associated with a weak silencer. We took IFITM3 as an example and did the same analysis as MMP14 (Supplementary Fig. 4). Both the accessibility of the enhancer 3 and the expression of IFITM3 are mildly decreased as the silencer gets more accessible (Supplementary Fig. 4c-e). The antagonism between the enhancer and the silencer does make the expression of IFITM3 more variable (Supplementary Fig. 4f).

To compare these two different patterns, we would like to propose two models about adversarial regulation on the same gene: a switch model and a competitive model. The switch model is tightly associated with strong silencers, which turn off the enhancer and gene transcription simultaneously. As a result, the enhancers’ function is restricted in a convergent triangular zone. That is, the enhancers only function in the absence of an active silencer, and the enhancers’ activity converges to 0 with the activation of the silencer. The competitive model is involved with weak silencers and may have a relationship with competitive combination with the promoter. As a result, gene expression is highly variable and can be finely controlled in both positive and negative ways.

### Delineation of transcription initiation activity of distal regulatory elements

Enhancer RNAs (eRNAs) are RNA molecules that are transcribed from genomic enhancer regions^23^. The previous study shows that the level of enhancer RNA expression positively correlates with the level of mRNA synthesis at nearby genes^24^. To decipher element functions in the transcription aspect, we quantitatively analyzed the transcription level of distal regulatory elements by leveraging 5’ scRNA-seq. We observed strong enrichment of RNA signal at the center of distal ATAC peaks (Fig. 6a). Transcription level and open chromatin states are positively correlated at sample level and cell type level with a large proportion of elements open but not transcribed (Fig. 6b and Supplementary Fig. 5a). To identify transcribed cis-regulatory elements (tCREs), in other words, open chromatin region with transcription initiation activity, at the whole organ scale, we applied a strict cut-off to each sample and merged tCREs lists into a master list of 190,356 regions (Supplementary Fig. 5b, c and Supplementary Table 7).

**Fig. 6.**
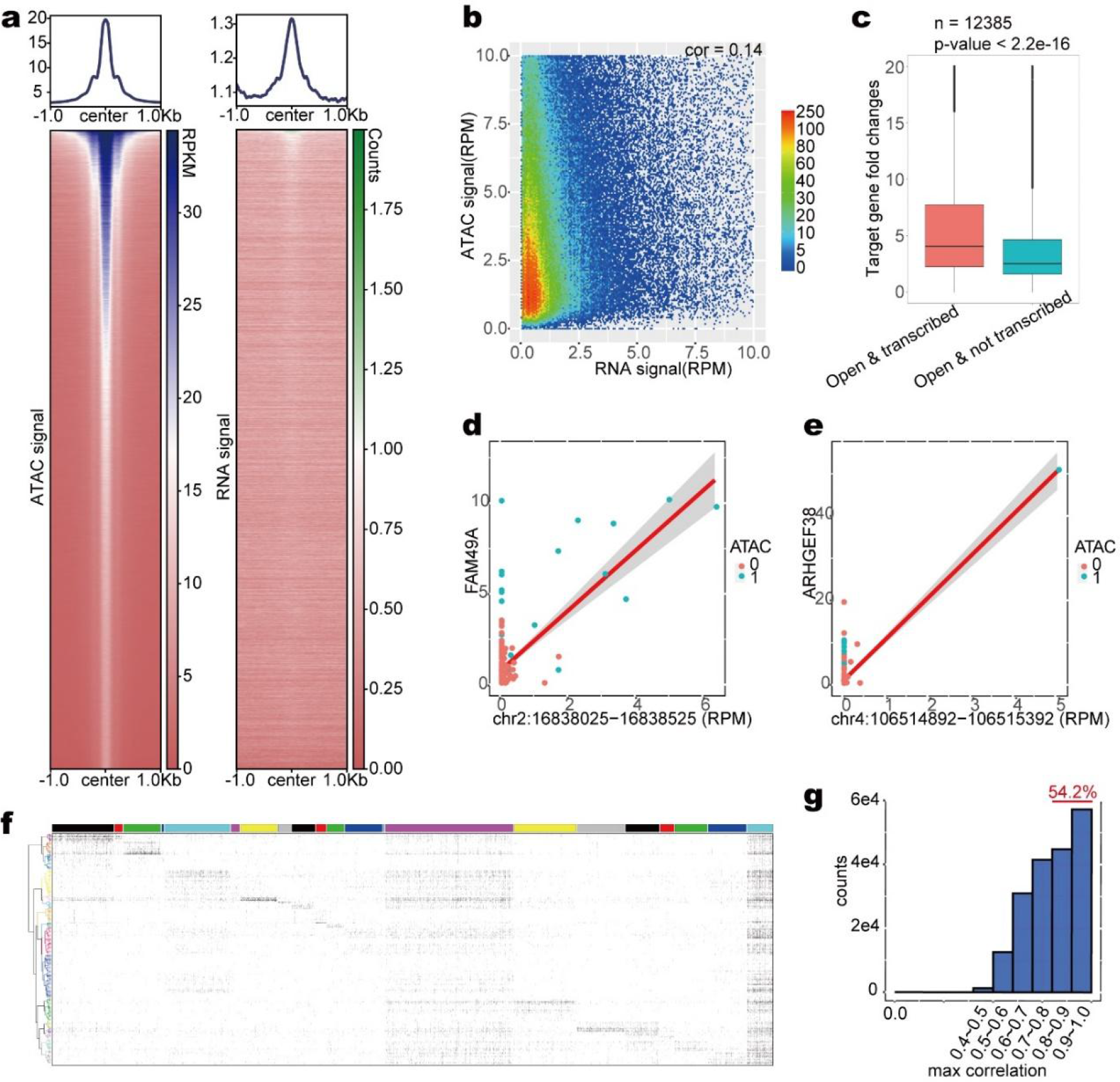
Transcription analysis uncover transcription-dependent and transcription-independent enhancers. a, Heatmaps showing the ATAC/RNA signal around distal ATAC peaks. (Left) ATAC signal of GW10 limb using RPM; (Right) RNA signal of GW10 Limb using number of read counts. b, Smooth scatter plot demonstrates the peak transcription level (x axis), along with ATAC signal intensity (y axis) of transcribed cis elements in each cell type. Only cis elements with open chromatin state are shown. c, Relationship between enhancer transcription and target gene expression. Boxes denote medians and interquartile ranges (IQRs, 25–75%), whiskers represent 1.5 x IQRs. d,e, Scatter plot demonstrates transcription level of the peak (x axis), along with transcription level of target gene (y axis) in each cell type. f, Transcription/not transcription at 190,356 transcribed cis elements (x axis) across 225 cell types (y axis). The color code on top represents 21 accessibility patterns. g, Frequency distribution of max correlation of co-expressed cis elements for each transcribed cis element.

For each cell type, about 10% of open chromatin regions have non-coding transcription start site signal on average. Combining tCREs with peak-to-gene links, we found cell types with transcribed enhancers have significantly higher expression levels of target gene than cell types with an un-transcribed enhancer (Fig. 6c). We further identified 1361 peak-to-gene links in an eRNAs-dependent manner, 206 of which were associated with TF-encoding genes^25^ (Supplementary Table 8). Open chromatin state is the necessary condition of transcription, and the level of eRNAs is a

determining factor in promoting target genes (Fig. 6d, e, and Supplementary Fig. 5f). To assess cell type specificity of the tCREs, we ordered tCREs according to their source peak groups and found a similar but more evident pattern with chromatin accessibility pattern, which may indicate higher specificity in cis-element transcription (Fig. 6f). We also note that universal open peaks have higher transcribed proportions and may have a specific function (Supplementary Fig. 5d). Multiple enhancers may be co-accessible and regulate the same gene. Based on this, we assumed that co-expressed cis-elements are likely to be functional elements instead of random non-coding transcription noise. We found about 54.2% of tCREs have a highly co-expressed patterner (cor>0.8) (Fig. 6g). We also found more than half of our defined enhancer-to-gene pairs are associated with un-transcribed cis-elements, most of which cooperate with another transcribed enhancer to regulate the same target gene (Supplementary Fig. 5e). What’s more, the remaining enhancers work alone without transcription signal, suggests that many enhancers function in a transcription-independent manner (Supplementary Fig. 5g). The precise molecular mechanism of different categories of enhancers needs further investigation.

### Enrichment analysis of GWAS signals in aCREs and tCREs LSFs

To further our understanding of lineage specifier families, we applied stratified linkage disequilibrium score regression^26, 27^ and evaluated heritability enrichment in 52 GWAS datasets (Supplementary Table 9) across these 20 LSFs. The spectrum of traits evaluated covered blood cell physical traits, neurological, immunological, gastroenterology, metabolomic traits from UK Biobank data^28^ and Broad LD Hub^29^.

We observed Immune-related LSF show similar heritability enrichment for immune traits (Supplementary Fig. 6a). Lupus, Crohn’s disease, Rheumatoid Arthritis are significantly correlated with immune-related LSFs (T-cells, Immune system, and macrophage). The strongest enrichment of heritability for immunoglobin A (IgA) deficiency is in T cells. Epithelial LSF dominated by different organs display specific enrichment features for organ-matched traits. Kidney epithelial are relevant with kidney−stone. The lung epithelial and gonad LSF both enrich in lung FEV1/FVC ratio. Likewise, some blood cells’ physiology traits and immune-related traits are significantly enriched in Erythroid LSF, T2D, and Fasting Glucose are highly correlated with Endocrine systems, which are consistent with prior knowledge.

Furthermore, we found that the enrichment tendency of heritability of two neuron LSFs is different. Neuron1 LSF, which is mainly contributed by the retina or neural portion of the eye, is part of the central nervous system. Neuron2 LSF, which is called enteric nervous system (ENS) LSF. The results of this GWAS heritability analysis showcase, several psychiatric traits, and major neurodegenerative disorders, like Schizophrenia, Neuroticism, highly correlate with Neuron1 LSF, in stark contrast with weak signal in Neuron2 LSF. It suggested that the eye is a ’window’ into the brain, the accessibility and organization of the retina make it a convenient research tool with which to study processes in the CNS^30^. Unexpectedly, the Eye-related open chromatin enriched variants of the muscle-skeletal system and connective diseases, which may suggest some unrecognized link.

The previous study suggests that some distal aCREs marked by ATAC-seq or DHS-seq signal don’t have enhancer activity. Those regions maybe not binding by TFs or not interact with the promoter to drive gene expression, even they are open. Meanwhile, those open chromatin regions which have transcription initiation activity (tCREs) are more likely to be active enhancers, since the RNA signal suggests they are accessible by Pol II. Thus, we wonder whether tCREs are more enriched with GWAS signals and functionally relevant. For each open chromatin LSF, we identify the corresponding tCRE LSF (Supplementary Table 9). We calculate the GWAS signals enrichment similar to aCREs as described above. Interestingly, we found the enrichment of some traits and disease related SNPs are higher in tCREs than in aCREs LSF (Supplementary Fig. 6b-d). To avoid the trait heritability difference is caused by captured SNP number from aCREs LSF and tCREs LSF. We calculate 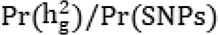 to measure LSF genetic associations and heritability. For Thyroid Disease, heritability was markedly enriched specifically within T cells associated tCREs LSF compared with aCREs, it indicated tCREs can capture trait heritability better than aCREs, it may cover more vital genetic signals (Supplementary Fig. 6b).

### Heritability enrichment identifies traits and disease-relevant fetal cell types

Many common diseases have a developmental origin. Despite the remarkable success of genetic signal mapping in GWAS, the functional interpretation of GWAS remains challenging. First, it is unclear in which tissues and cell types these variants are active, and how they disrupt specific biological networks to impact disease risk. Second, most disease-associated variants are located in non-protein-coding regions of the genome, and many are far away from the nearest known gene. We have evaluated the genetic risk of traits and disease for LSF, however, the most relevant cell types of certain diseases during organogenesis are poorly understood. CREs are bits of noncoding DNA that regulate the transcription of nearby genes. Here we can use each cell type top 10,000 specific CREs^9^ to explore the cellular context in which disease-associated variants act.

The results revealed that risk variants for kidney stones and chronic kidney diseases were enriched in kidney tubule cells (Fig. 7a). For tubule cells, it comprised distinct subpopulations with differentially accessible chromatin regions. We further provide a finer genetic signal map of the tubule subpopulation. Distal tubule cell shows higher enrichment (q value <0.05) for kidney function-related traits (eGFR, BUNM, Urate) from the study by Wuttke et al. and Teumer et al^31, 32^, and S-Shaped body cell type and LoH cells are both relevant to the kidney stone. Likewise, we find endocrine cells, which showed significant enrichment for fasting glucose (Fig. 7b).

**Fig. 7.**
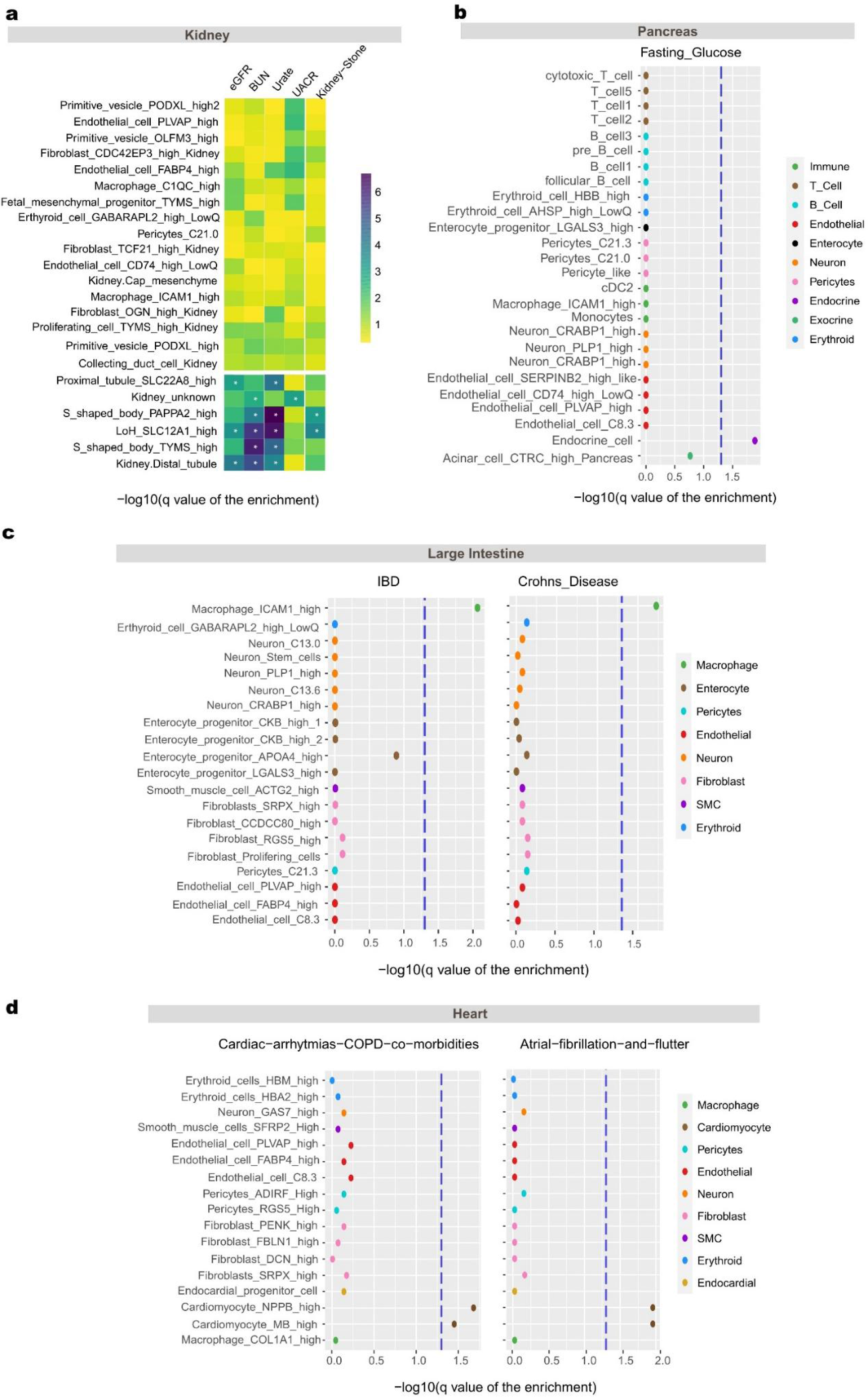
Enrichment analysis of GWAS signals in cell-type-specific chromatin regions. a, S-LDSC results suggests these disease and traits’ susceptibility and heritability are cell type specificity. Heatmap show cell-type-specific enrichments of the heritability signal for kidney stone and CKD diseases in kidney tissue, significance level (q<0.05) are indicated with an asterisk. b, Dot plot show cell-type-specific enrichments of the heritability signal (y axis) for diabetes in pancreas tissue, the blue dotted line indicates significant threshold (q value of 0.05). c, Dot plot show the cell-type-specific enrichments of the heritability signal for two typical inflammatory bowel diseases across all cell types in large intestine. d, Dot plot show cell-type-specific enrichments of the heritability signal for heart traits in heart tissue.

Dot plot shows the -log10(q value of enrichment) for two chronic Inflammatory bowel diseases (IBD, Crohns’ Disease) across all cell types in the large intestine (Fig. 7c). Only one digestive-system sourced macrophage has significant enrichment. It consisted of a recent study that reported a subtype of NOD2-driven Crohn’ s disease leads to dysregulated homeostasis of activated fibroblasts and macrophages^33^.

The most relevant cell type of heart traits cardiac arrhythmias and atrial fibrillation and flutter (AF) and Cardiac arrhythmias COPD comorbidities are cardiomyocytes (Fig. 7d, and Supplementary Fig. 6e). AF risk variant (rs7789585) is located in a cardiomyocyte’s specific open chromatin region, which resides in the second intron of the KCNH2. Co-accessibility analysis suggests that KCNH2 is likely the target. This observation is consistent with a recent report that cardiomyocyte enhancers of potassium channel gene KCNH2 may be affected by noncoding risk variants associated with AF^34^. Collectively, we have assigned the most relevant fetal cell type for 10 traits or diseases (Supplementary Table 9).

### Cell type of origin for cancer

Cells from fetal tissue and tumor both grow and divide rapidly, and they share common cell surface markers and oncofetal antigens, include carcinoembryonic antigen (CEA), alpha-fetoprotein (AFP)^35^. To a certain extent, malignant tumor regulatory mechanisms resemble fetal cells, the fetal tissue in a single-cell resolution may provide the answer of the cell type of origin for the tumor. For example, a recent study has found that most adrenal NB tumor cells transcriptionally mirror early human embryos’ noradrenergic chromaffin cells^36^. Moreover, another recent study reported a shared immunosuppressive oncofetal ecosystem in fetal liver and hepatocellular carcinoma^37^, suggesting fetal tissue may provide a better understanding of the tumor ecosystem. The large-scale cross organ datasets generated in our study allow us to explore the similarity of fetal cell types with multiple cancer types. To ensure accuracy, we pay more attention to 9 tumor types from TCGA which have corresponding fetal tissue in our datasets, and their chromatin state was profiled by bulk ATAC-seq in a previous study^38^.

For each tumor sample, we inferred the putative cell type of origin based on the chromatin accessibility similarity with fetal cell types using Jaccard distance (Supplementary Table 10). We observed almost all patients show accordant preference on specific cell types based on chromatin accessibility and found cancer-associated cell types. Across 41 stomach adenocarcinoma (STAD) samples, the fetal stomach cell types which show the highest similarity score consistently to be Surface Mucous Pit Progenitor cells (Fig. 8b, d), which make mucus and stomach juices.

**Fig. 8.**
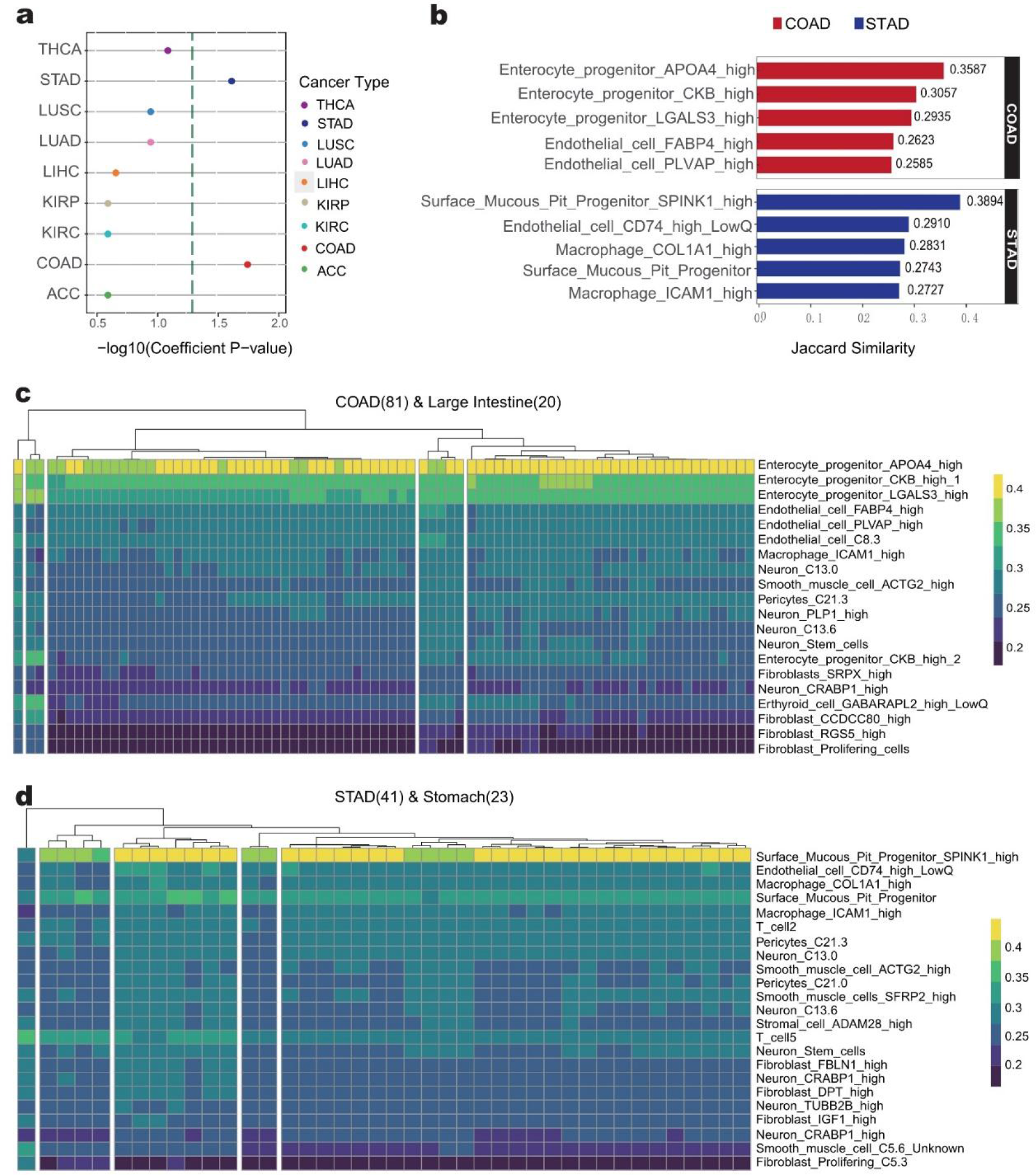
Link fetal cell type with cancer at chromatin stat level. a, Rank of tumor type relevance to proliferative state cell types based on hypergeometric test. x-axis is the -log10(p value), blue dotted line is p value of 0.05. b, Bar plot showing the Jaccard similarity Score of top5 similar fetal cell types for Colon Adenocarcinoma (COAD) and Stomach Adenocarcinoma (STAD). c, Jaccard similarities of chromatin state from 81 Colon Adenocarcinoma individuals (y axis) with the cis-elements of 20 cell types in large intestine (x axis). d, Jaccard similarities of chromatin state from 41 Stomach Adenocarcinoma (STAD) individuals (y axis) with the cis-elements of 23 cell types in stomach (x axis).

In a similar fashion to previous analysis of STAD-associated fetal cell types, we summary the top5 most similar fetal cell types for each tumor (Fig. 8b, and Supplementary Fig. 7a). Meanwhile, 54 cell types in our data have been annotated as a proliferative state based on CytoTRACE^39^ inference and unique gene expression (Supplementary Table 10). To investigate whether the cancer-associated fetal cell types are enriched in proliferate or progenitor cell types, we use a hypergeometric test to compute the statistical significance of the intersection of cancer associated cell types and proliferative state cells. We found colon adenocarcinoma (COAD) and STAD are clearly different from the other 7 cancer types, their associated fetal cell types are significantly enriched proliferate states (Fig. 8a). The COAD-associated cell type is enterocyte progenitor cell that sustains proliferating state in the large intestine, while STAD associated cell type is surface Mucous Pit progenitor in the stomach (Fig. 8c, d). Moreover, both of them show similar chromatin states nearby CEA family genes with cancer (Supplementary Fig. 7b, c).

## DISCUSSION

In this study, we leveraged single-cell profiling of RNA and chromatin to perform integration analysis and construct cis-regulatory elements atlas. The scale of the current analysis helped us to discern more details on the biological phenomenon and better understand transcription regulation. By comparing with the VISTA database, we got to know validated enhancers are open in which cell types. By integrating motif enrichment and gene expression, we confirmed transcription factors acting as key regulators of dynamics of open chromatin and lineage differentiation. By combining positive cis-elements with negative cis-elements, we found mutually exclusive modules regulating the same genes in a cell-type-specific manner, which may provide a potential way for disease treatment.

The cis-regulatory elements atlas of the current study provides a snapshot of fetal development. It would be more valuable to sample in continuous stages, offering a spatiotemporal perspective of lineage hierarchy and transcription regulation. More advanced experimental technologies and algorithms will emerge and set a foundation for better resolving fetal development sometime in the future.

Fetal tissue with persistent differentiation potential finally developed functional mature normal adult tissue, whereas it also can switch to tumor disordered proliferation state in the oncogenic mutations stimulate. It looks like a one-direction irreversible event, whereas tumor tissue can break this order, it reactivates some cis-elements with normal fetal tissue which keep silent in adult tissue, switch cell status to benefit tumorigenesis. Our study builds a bridge between the two physiological states based on the similarity of the open state of chromatin and provides a new perspective for the exploration of the developmental origin of tumors. We systematically summarized fetal cell types have a similar regulatory mechanism with 9 primary tumors. In addition, for TCGA bulk level ATAC-Seq data of tumor tissues, it can observe cellular composition heterogeneity and complex microenvironment in tumor samples. And our findings show these oncofetal antigens are cell type-specific open in fetal tissues, which prefer proliferating state cell types with persistent multilineage differentiation potential, and these genes are also reactivated in tumor cells, which seems to support the previous hypothesis. However, we haven’t detected adult tissue and cancer tumor chromatin state at a single cell level, so, we can’t verify whether these cell types truly can happen oncofetal reprogramming.

## Acknowledgments

Funding: This work was supported by the Strategic Priority Research Program XDB38020500 to L.J., National Key R&D Program of China 2019YFA0801702 to L.J., the Natural Science Foundation of China 31970760 to L.J.

## Author contributions

L.J., J.L. and P.P. conceived the study. H.Y. and Y.K. facilitated its designs. P.P., Y.K., S.J., and Y.L collected embryos sample. Y.K., S.J., T.Z., and Y.L performed scRNA-seq and scATAC-seq library construction. H.Y. and N.A. performed the bioinformatics analyses. N.A., H.Y., Y.K., X.C., L.J., and J.L. interpreted the data. H.Y., N.A., J.L. and L.J. wrote the paper with the assistance of the other authors.

## DATA AND CODE AVAILABILITY

The accession number for the sequencing data reported in this paper is submitted to Genome Sequence Archive for Human (GSA-Human): https://ngdc.cncb.ac.cn/gsa-human/s/x70211Pp. The processed files are uploaded to Figshare: https://figshare.com/projects/HumanProject/122983. All codes are available upon reasonable request.

## Supplementary information

Legends for Supplementary Files

**File S1** | Metadata of cells in scRNA-seq data. Includes sample metadata, per-cell QC stats, cluster id and cell type annotation.

**File S2** | Metadata of cells in scATAC-seq data. Includes sample metadata, cell type annotation and various Cell_ID information for each software.

**File S3** | Gene count matrix of cells in scRNA-seq data in RDS format. Includes expression UMI values for each gene in each cell.

**File S4** | Peak count matrix of cells in scATAC-seq data in RDS format. Includes insertion counts within each peak in each cell, while the maximum value was set to 4.

**File S5** | Normalized peak by cell type matrix in RDS format. Includes normalized peak accessible values (reads per million reads/100) for each cell type.

**File S6** | Binary peak by cell type matrix in RDS format. Includes binary values for each cell type, where 1 denotes accessible and 0 denotes inaccessible.

**File S7** | Seurat object of 185,061 high-quality cells in scRNA-seq data. Includes count matrix, low-dimension embedding and cell informations from the global perspective.

**File S8** | Seurat object of average profiles of 335 cell types in scRNA-seq data.

**File S9** | tCRE transcription intensity matrix of each cell type in scRNA-seq data in RDS format. Includes RPM value for each peak in each cell type.

**File S10** | 3D animated scatter plot representing relationship between gene expression level and enhancer/silencer activity in gif format. Pattern 1 is related to Fig. 5g, while pattern 2 is related to Supplementary Fig. 4f.

The Supplementary Table S1 can be downloaded from https://figshare.com/ndownloader/files/30790600

The Supplementary Table S2 can be downloaded from https://figshare.com/ndownloader/files/30790603

The Supplementary Table S3 can be downloaded from https://figshare.com/ndownloader/files/30790606

The Supplementary Table S4 can be downloaded from https://figshare.com/ndownloader/files/30790609

The Supplementary Table S5 can be downloaded from https://figshare.com/ndownloader/files/30790612

The Supplementary Table S6 can be downloaded from https://figshare.com/ndownloader/files/30790615

The Supplementary Table S7 can be downloaded from https://figshare.com/ndownloader/files/30790588

The Supplementary Table S8 can be downloaded from https://figshare.com/ndownloader/files/30790591

The Supplementary Table S9 can be downloaded from https://figshare.com/ndownloader/files/30790594

The Supplementary Table S10 can be downloaded from https://figshare.com/ndownloader/files/30790597

These processed files are also uploaded to Open Archive for Miscellaneous Data (OMIX) database: http://ngdc.cncb.ac.cn/omix/preview/MCawh0yL.

## METHOD DETAILS

### Tissue acquisition and processing

The study of human embryos was approved by the Reproductive Study Ethics Committee in Peking Union Medical College Hospital, Beijing, China. All tissue samples used for this study were obtained with written informed consent from all participants. Samples from surgically removed aborted fetal tissues were collected into Leibovitz’s L-15 (11415064, Gibco) plus with 10% fetal bovine serum (FBS) right after resection and immediately transported on ice from hospital to the laboratory in less than 1 h.

We collected 4 individual ranging from: 6 PCW (post conception weeks), 10 PCW to 16 PCW and a total of 28 samples (15 organs or tissues): spleen, pancreas, liver, thymus, thyroid, lung, stomach, small intestine, big intestine, kidney, male gonad, female gonad, fore-limb, heart, and eye were including (Supplementary Table S1).

Each organ was dissected and washed with DPBS twice, then collected in 1.5 mL EP tubes.

### Single cell preparation and Nuclei Isolation

Tissues were minced into pieces (∼1 mm) on ice using scissors, and digested into single-cell suspensions with 1 mg/ml type II collagenase (17101015, GIBCO) and 1 mg/ml type IV collagenase (17104019, GIBCO) for 30min at 37 ℃ with intermittent shaking. The dissociated cells were separated and remaining undigested tissue were digested again with fresh digestion buffer. Digested suspension was passed through 70um strainer (Biologix).

Dissociated cells were centrifuged at 300 g for 5 min at 4 °C, then re-suspended in 1 mL of cold DPBS with 0.1% BSA. After passing through a 40um cell strainer (Biologix), cells were washed twice, centrifuged at 300 g for 5 min at 4 °C, re-suspended in cold DPBS with 0.1% BSA at a density of 1×105 cells/ml, and stored on ice before scRNA-Seq and nuclei isolation.

To isolate nuclei, the half of the cell pellets were re-suspended in 100 uL chilled lysis buffer (10 mM Tris-HCl, pH 7.4, 10 mM NaCl, 3 mM MgCl2, 0.1% NP40, 0.1% Tween-20, and 0.01% digitonin - from ^1^ supplemented with 1% BSA), and pipette mix 10X. After incubation for 5 min on ice, add 1 ml chilled Wash Buffer ((10 mM Tris-HCl, pH 7.4, 10 mM NaCl, 3 mM MgCl2, 0.1% Tween-20, 1% BSA) to the lysed cells. Pipette mix 5x, then centrifuged at 300 g for 5 min at 4 °C. Based on number of cells used for isolation and assuming ∼50% nuclei loss during cell lysis, resuspend in chilled Diluted Nuclei Buffer (PN-2000153, 10x Genomics). If cell debris and large clumps are observed, pass through a cell strainer. For low volume, use a 40 µm Flowmi Cell Strainer (H13680-0040, Bel-Art) to minimize volume loss.

### sc-RNA-seq Libraries Construction and sequencing

Single cell RNA-seq was performed using the Single Cell 5’ RNA Reagent Kits (10x Genomics, Pleasanton, California) according to the manufacturer’s instruction. The aimed target cell recovery for each library was ∼9,000 cell per sample. In brief, cellular suspensions were loaded on the sample chip in the Chromium Controller instrument (10X Genomics) to generate single-cell Gel Bead-In-Emulsions (GEMs). GEM-reverse transcription (RT) was performed in a Veriti 96-well thermal cycler (BioRad, 1851197). After RT, GEMs were harvested and the cDNAs were amplified, and cleaned up with SPRIselect Reagent Kit (Beckman Coulter, Pasadena, CA).

Indexed sequencing libraries were constructed using Chromium Single-Cell 3′ Library Kit or Single Cell 5’ Library kit based for enzymatic fragmentation, end-repair, A-tailing, adaptor ligation, ligation cleanup, sample index PCR, and PCR cleanup. Libraries were quantified using Bioanalyzer (Agilent) and QuBit (Thermofisher) analysis and then sequenced in NovaSeq 6000 (Illumina, San Diego, CA) with a 150-bp paired-end read length, targeting a depth of 50,000–100,000 reads per cell.

### sc-ATAC-seq Libraries preparation and sequencing

The scATAC library was prepared using the 10x Genomics platform with the Chromium Single Cell ATAC Library & Gel Bead Kit (10x Genomics, Pleasanton, California) as instructed by the manufacturer. A total of 15,000 nuclei per sample were used as input for single-cell ATAC-seq following the manufacturer’s instructions. Briefly, after tagmentation, the cells were loaded on a Chromium Controller Single-Cell instrument to generate single-cell Gel Bead-In-Emulsions (GEMs) followed by linear PCR as described in the 10X scATAC-seq protocol using a Veriti 96-well thermal cycler (BioRad, 1851197). After breaking the GEMs, the barcoded tagmented DNA was purified with SPRIselect Reagent Kit (Beckman Coulter, Pasadena, CA) and further amplified to enable sample indexing and enrichment of scATAC-seq libraries. The final libraries were quantified using Bioanalyzer (Agilent) and QuBit (Thermofisher) analysis and then sequenced in Nextseq 550AR or NovaSeq 6000 (Illumina, San Diego, CA) with a 50-bp paired-end read length, or MGISeq-2000FCL (MGI Tech Co., Ltd., China) with 100-bp paired-end read length targeting a depth of 30,000–50,000 reads per cell.

### scRNA-seq Data processing

FASTQ files generated from sequencing were used as inputs to the 10X Genomics Cellranger (3.1.0) RNA pipeline using default arguments. Briefly, de-multiplexed reads were mapped to the hg19 genome by STAR. Filtered feature-barcode matrix containing feature, barcode list and matrix was generated and as input to Seurat (version 3.2.3).Cells with low complexity(fewer than 400 expressed genes) were excluded; cells with mitochondrial read fraction outside 10 percent were also cleared out. The Seurat (version 3.2.3) workflow were run separately on each sample, most of these parameters have default setting,and the resulting files were used for further processing. Doublet was estimated for each 10x sample by applying the ‘doubletFinder_v3’ function in the DoubletFinder package (version 2.0.2), which is implemented to interface with Seurat.This function predicts doublets according to each real cell’s proximity in gene expression space to artificial doublets created by averaging the transcriptional profile of randomly chosen cell pairs.

### scRNA-seq clustering and cell type annotation

Each dataset was integrated together using the ‘merge’ function in the Seurat package.High quality cells from all samples were merged and normalized (normalization.method = “LogNormalize”, scale.factor = 10000). Highly variable genes (HVGs) had significantly variance were retained (selection.method = “vst”, nfeatures =10000).Notably, we regressed out the difference between the G2M and S phase scores (vars.to.regress = S.Score - G2M.Score) to mitigate the effects of cell cycle heterogeneity in scRNA-seq data. Next ,batch effects were removed by harmony on 75 principal components computed from the HVGs only. Correction was performed between the samples of each time point, this method was carried out on the whole atlas dataset, and Harmony embeddings calculated from this batch-corrected principal component analysis were used for all further analysis steps.We used shared-nearest-neighbours (SNN) and Louvain method to cluster cells and identified 42 distinct major clusters (dims = 1:75 and resolution = 0.3). To identify finer substructure from these major clusters, each cluster underwent a second round of clustering using the same methods as above with resolution range from 0.2 to 0.6, respectively. We further remove 3192 cells from 21 sub clusters with doublet ratio previous calculated higher than 55%. Finally, we identified a total of 331 sub clusters. Differential expression analysis for each cluster was performed by using the “FindAllMarkers” function with default Wilcoxon rank-sum test. Cell types were assigned to each sub cluster based on the enrichment of cell type of Human Cell Landscape (HCL) and the expression of known marker genes. Details of cell type annotation information are listed in Supplementary Tables 2.

### scATAC-seq Data processing

After sequencing, FASTQ files were processed with 10X Genomics Cellranger-atac (1.2.0) pipeline with default parameters. Briefly, the reads were aligned to hg19 using BWA to generate fragment files. Only fragments with MAPQ > 30 on both reads were retained. Each unique fragment is associated with a single cell barcode. After filtering low quality barcodes and removing PCR duplicates, a total of ∼3.1 billion read pairs were retained from scATAC-seq. These reads constitute 269,920 valid cells. The output HTML files containing metrics and library information are organized into a table (Table S1). The output fragment files were loaded into ArchR to generate cell-bin matrix. Briefly, we exclude low-quality cell barcode based on loose quality control parameters: 200 unique fragments per cell and a transcription start site (TSS) enrichment score of 4. Then, we used computational framework bap (bead based ATAC processing) to combine cells which have similar fragments but with different barcodes. New fragment files generated by bap2 were loaded into ArchR again. We picked the top 12,000 cells with the highest TSS enrichment score to remove the effects of cell numbers per organ and adopted a strict quality control parameter: 1000 unique fragments per cell. Finally, we filtered the doublets with addDoubletScores function in ArchR and attained final cell-bin matrix for further analysis. Finally, 230,732 high-quality cells with balanced sample sources are used for downstream analysis.

### Cell type identity assignment of scATAC-seq data

To annotate cell types for scATAC-seq data, we transferred cell type labels from scRNA-seq to scATAC-seq data within paired assays. First of all, we arranged 331 cell types by transcriptomic similarity and pre-divided them into 65 groups by using the R package dendextend. Then, we performed two rounds of label transferring using ArchR, which utilize Seurat’s canonical correlation analysis (CCA) based integration infrastructure. For the first round, we transfer 65 cell type group labels with unconstrained integration mode. For the second round, we transferred 331 cell type labels with constrained integration mode. Briefly, dimensionality reduction of whole scATAC-seq dataset was performed by using Latent Semantic Indexing (LSI). Cells were clustered by Louvain algorithm with r=7 (seurat’s FindClusters) and visualized by UMAP. Through first-turn label transferring, we identified which cell type group labels from the scRNA-seq data are most abundant in each of scATAC-seq clusters. We constructed a “groupList” which contains 65 pair of lists of cell IDs across scRNA-seq and scATAC-seq dataset. Then we pass this list to the ‘groupList’ parameter of the ‘addGeneIntegrationMatrix()’ function in ArchR and performed second-turn label transferring constrained in each group and sample. We achieved a median prediction score of 0.58-1.0 across 28 samples. 283 cell types were successfully transferred. The cell types with cell number higher than 50 were performed peak calling by using macs2. Totally, 848,475 non-overlapped 501bp fixed-width master peaks was generated. Any peak that directly overlaps with most significant peak was removed. After filtering cell types with less than 50 cells or 20,000 peaks, we got 225 cell types with paired pseudo-bulk profiles of gene expression and chromatin accessibility.

### Genome browser visualization of two assays

Firstly, we used samtools to merge sample bam files together. Secondly, we used filterbarcodes command in the Python package sinto (v0.1, https://github.com/timoast/sinto) to get bam file for each cell type. Finally, we generated bigWig files using bamCoverage program in Deeptools2 with parameter “-noralizeUsingRPKM’’ and visualized them in IGV (version 2.8.13) (Fig 1D).

### Generate DNA accessibility patterns using binary peak-by-cell type matrix

We constructed a binary matrix Mp2ct consisting of the presence or absence calls of the master peak list (n = 848,475) across 225 cell types. Mp2ct (225*848,475) was clustered by rows and columns separately. Firstly, we selected top 200,000 most variable peaks across cell types as features. Secondly, we calculated distance between each cell type using (1-pearson correlation). Thirdly, we did hierarchical clustering using calculated distance using ward.D2 algorithm (Fig 1C). For column clustering, we unitized 2-norm of each column of Mp2ct to 1 and got a normalized matrix Mnor. Then we took cell types as features and applied K-means to 848,475 columns of Mnor in Hartigan-Wong algorithm. We tested different K according to an arithmetical sequence, and selected satisfactory one (K = 21) based on internal structure of Mp2ct heatmap organized in clustering results. Lastly, we manually adjusted peak group orders to visualize the binary matrix in a fashion of neatly arranged blocks on the diagonal. Note that the same procedures were also applied to identify sub patterns of cell types of kidney epithelial.

### Overlap of the ATAC peaks with consensus human DHSs

To assess the overlap between our ATAC peaks and DHSs from large-scale bulk DNase-seq, we obtained index of consensus human DHSs from ENCODE Project and computed intersection as well as subtraction between two datasets. The comparison were made in two cases: whole dataset level (Fig S2A); among corresponding primary tissues (Fig 2B). To explore differences between datasets in case two, we also calculated tissues/cell types contributions to datasets specific peaks (Fig 2C and S2B). Note that one peak may be calculated repeatedly, but only a limited overlap exists between sub cell types (Fig S2C). Lastly, two-tailed Student’s t test was conducted between contributions from common cell types and contributions from organ specific cell types (Fig 2D).

### Enrichment analyses for enhancers from the VISTA enhancer database

VISTA validated elements were downloaded from https://enhancer.lbl.gov on 27 September 2020. To attain the expression pattern of each enhancer, we used advanced search on the website and downloaded the enhancers from corresponding organs (eye, heart, limb, liver and pancreas) in turn. Firstly, a global comparison was made regardless of organ source (Fig S2D). Secondly, we characterized accessibility pattern of enhancers across different cell types using binary matrix (Fig S2E). Finally, we shuffled organ peaks 3 times as background for each test, and calculated observed to expected (median value of overlaped peaks in random situation) ratio as enrichment to eliminate quantity effects (Fig 2E). We repeated the above operation and got enrichment in cell type level (Fig 2F).

### Transcription factor motif enrichment and expression analysis

The findMotifsGenome.pl in HOMER was used to calculate TF motif enrichments in different peak groups (Fig 2A and 3A) with parameter “-size 400”. Only the top 10 motifs of each peak groups were selected to perform visualization and annotate peak groups. Gene expression levels of TFs were normalized across cell types by Z-score and visualized using ‘DotPlot()’ function in Seurat. Note that a gap exists between TF names from HOMER and official gene symbols. We filled the gap by taking two strategies: convert lower-case characters to upper-case to see if matching any official gene symbol; manually search the TF names on GeneCards database to see if matching any aliases of a gene. An organized csv file was available on the website.

### Finding Instance of Specific Motifs

To recover the locations of each motif found in the motif discovery process, we ran the findMotifsGenome.pl again with parameter: -find SIX2.motif. The recovered peaks were defined as TF target peaks.

### Linking regulatory elements to cognate genes

By ArchR, we leveraged the gene expression data and created a correlation-based map between chromatin accessibility peaks and their cognate genes directly. Briefly, an approach introduced by Cicero is adopted to create low-overlapping aggregates of single-cell profiles. Aggregates with greater than 80% overlap with any other aggregate are filtered in order to reduce bias. Then we leveraged scATAC-seq data and integrated scRNA-seq data to look for correlations between peak accessibility and gene expression. These putative gene regulatory interactions were predicted using the “ getPeak2GeneLinks ” function with default parameters in ArchR. We searched a region of ± 250kb for each gene and filtered peaks which were proximal to TSS (± 1kb). Links with absolute value of correlation larger than 0.45 or less than -0.40 were used for downstream analysis. Positive links are defined as enhancer-gene links, and negative links are defined as silencer-gene links.

We repeated these procedure in whole organism level as well as within each organ. To retain reliable linkages against random noise, we filtered links that only shows in one condition and merged the leftovers into 155,620 positive peak-to-gene links (associated with 108,699 peaks and 12,783 genes) and 34,287 negative peak-to-gene links (associated with 23,392 peaks and 7,628 genes).

### Association with ReSE-identified silencers

To validate our data, we did overlap between correlation-based silencers and ReSE-identified silencers by using intersectBed. Then, we applied the same correlation-based methods linking ReSE-identified silencers to cognate genes. Only the negative correlated links were taken into consideration. We set the region as ±500kb for each silencer, and assigned the gene with the smallest correlation to this silencer. 2113 of 5472 ReSE-identified silencers were assigned with a target gene.

### Classification of silencers

To determine the class of each sliencer, we focus on the distribution of gene expression with different peak accessibility and see if there is a sharp decline once the peak accessibility reach a critical value.

For each silencer, we simply take a list of value of 1/10, 2/10, …, 8/10*Max, where Max denotes the max value of the peak accessibility. For each value i, we seperate cell types into two group, one with peak accessibility more than i (group i1), and one with peak accessibility no more than i (group i2). If either of the groups has less than 6 cell types, we skip the value i. Then we calculate mean value and variance for each group (Ei1, Vari1 for group i1; Ei2, Vari2 for group i2). A silencer is classfied as strong silencer only if Ei2/ Ei1 > 3 and Vari2/ Vari1 > 3 for any of the value i. We tested the classifier on both correlation-based silencers and ReSE-identified silencers, and got the same result with the independent man-made result.

### Trajectory inference with Palantir

The Palantir workflow consists of three core steps to align cells along differentiation trajectories. Palantir also includes visualization tools to help explore trajectories and capture the stochasticity in cell fate determination.

Dimensionality reduction with force-directed layouts (FDL). Firstly, we exported cell-peak matrix and cell-gene matrix from ArchR and tranferred it into mtx format. Secondly, the matrices were loaded into Palantir via ‘scanpy.read_10x_mtx()’ function. To settle the high sparsity of scATAC-seq data, we searched 50 nearest neighbors for each cell via ‘scanpy.pp.neighbors’ function, and aggregated single-cell profiles using following formula:

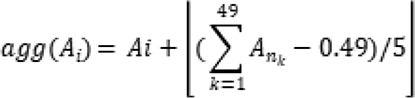

where *Ai* denotes count number of cell *i* on a peak and *n_k_*(*k*=1… 49) denotes neighborhood cells. Thirdly, the aggregated ATAC profiles were used for FDL visualization via ‘harmony.plot.force_directed_layout()’ function.

Integration with scRNA-seq data. To integrate transcriptome into the Palantir framework, we took the diffusion maps of the scATAC-seq data from ‘palantir.utils.run_diffusion_maps()’ function. Using the same diffusion maps, we can visualize gene expression levels on the same FDL plot. Then we plotted maker genes on FDL to attain cell type locations.

Grouping cells into different trajectories. We first specifying an approxiate early cell and terminal cells based on marker genes. Next we ran Palantir core function on scATAC-seq data by ‘palantir.core.run_palantir()’. Palantir generates the following results: pseudo time ordering of each cell; terminal state probabilities of each cell; a quantiative measure of the differentiation potential of each cell. We partitioned cells into trunk and branches according to terminal state probabilities. Cells with balanced probabilities are defined as trunk and are used for start of lineage differentiation. Pseudo time from 0 to 1 is used to order cells.

### Characterization of TF related enhancer elements and genes along differentiation

We characterized chromatin accessibility of TF related enhancer elements and expression level of TF related genes by using Locally Weighted Linear Regression (Loess). Briefly, we extracted profiles of chromatin accessibility/gene expression from cell-peak/cell-gene matrix and ordered cells according to pseudo time. We truncated the top 5% and bottom 5% among all cells and applied Min-Max normalization to each profile to make cross-data comparison. Finally, each profile of chromatin accessibility/gene expression combined with pseudo time was fitted with Loess model by ‘geom_smooth()’ function.

### Generating paired DNA accessibility patterns and gene expression patterns

To visualize DNA accessibility patterns and gene expression patterns, we firstly calculated average gene expression levels/DNA accessibility for each cell type. For scRNA-seq data, we used ‘Seurat::AverageExpression()’ function to average gene expression by cell types. For scATAC-seq data, the read count of each cell in the cell-peak matrix was normalized to 10,000. All cells with the same cell type label were pooled together to get the average DNA accessibility. Then we took enhancers/silencers related to the same gene as an unit, and used average value to represent accessibility of enhancers/silencers to the gene. Next we drew heatmap for gene expression patterns of 108 genes identified above, and clustered genes using R package ComplexHeatmap with parameter: cluster_columns = T. Enhancers and silencers are in the same order as their linked genes, and were visualized with heatmaps. Gene expression levels was normalized across cell types in Z-score and limited from -2 to 2 for the visualization. DNA accessibility was normalized across cell types in Z-score and limited from 0 to 2 for the visualization.

### Colocalization of scATAC-seq signal and 5’ scRNA-seq signal

To distinguish transcription at CRE from mRNAs, we firstly filtered scRNA-seq reads proximal to TSS (±1kb) or overlaped with any exon. We have 8 samples with paired-end sequencing and 18 samples with single-end sequencing on read 2 (median fragment size: 350bp). To uncover transcription start sites precisely, we focused on read 1 for paired-end sequencing, and shifted upstream 200bp for single-end sequencing. Only the very beginning 50bp of each read are used for downstream analysis. We calculated scATAC-seq signal and 5’ scRNA-seq signal per distal ATAC peak and prepared an intermediate file via ‘computeMatrix’ in deeptools (version 3.3.0). Finally, we visualized all the results in paired heatmaps via ‘plotHeatmap’ (Fig 5A).

### Identifying transcribed cis-regulatory elements

To identify transcribed cis-regulatory elements, we started from sample levels and chose representative characteristics. For each sample, transcribed cis-regulatory elements are defined as open and significant transcribed. We used ATAC data to call peaks via macs2. Then we calculated local RNA siganl enrichment by using the ratio between core read count and average background read counts ( (upstream 500bp + downstream 500bp)/2 ). Only the peaks with more than 5 read count and more than 1.5 local RNA siganl enrichment are considered as significant transcribed. Finally, we merged transcribed cis elements from each sample into a master list of 190,356 peaks. For each cell type, any of the 190,356 peaks with open state and read count larger than 3 are considered as transcribed cis elements.

### Identifying Cell type Specific aCREs

In each organ, we calculate specificity score for every cell type based on the cells versus 84K aCREs matrix by ‘Specificity scores’ preprint protocol V1.01^2, 3^ which provided by Silvia et al. Then rank theses aCREs based on the specificity score, the top 10,000 most specific CREs per cell type is used in downstream analysis.

### Enrichment analysis of Heritability

Partitioned heritability was measured using LD Score Regression v1.0.0^4, 5^ to identify enrichment of GWAS summary statistics among lineage specifier families (LSF). To do so, first all necessary data set needed to run S-LDSC including baseline scores, PLINK files, frequency files, weights, and SNPs, were downloaded from the Broad Institute. All files were ‘ 1000G_Phase3’ versions (See TableS6). Additionally, Roadmap Epigenetic Project LDSC files were used as additions to the baseline model as was done in a previous application of LDSC on ATAC seq data. We obtained GWAS summary statistics data from the UK Biobank project as processed by the Neale lab (http://www.nealelab.is/uk-biobank/). Summary statistics for 52 GWAS were obtained from have been processed into LDSC-format using the ‘munge_sumstats.py’ script.

Firstly, annotation file was created which marked all HapMap3 SNPs that fell within top 10K CREs for each cell type, which were ranked by cell type-specificity scores. Then LD-scores were calculated for these SNPs within 1 cM windows using the 1000 Genomes data with the ‘ldsc.py’ script. These LD-scores were included simultaneously with the baseline distributed annotation file from 1000 Genome project phase 3 with population code EUR and another baseline model from Roadmap Epigenetic Project LDSC files. Subsequently, the heritability explained by these annotated regions of the genome was assessed from these genome-wide association studies: The enrichment was calculated as the heritability explained for each phenotype within a given annotation divided by the proportion of SNPs in the genome and Benjamini–Hochberg FDR correction (Benjamini and Hochberg, 1995) was used to correct for multiple comparisons. Partitioned heritability calculations for all traits were combined and analyzed in R. The creation of plots was carried out using custom R scripts. The level of significance was set for LDSC results as the Bonferroni corrected P-value when take into account all summary statistics and cell populations tested.

Heritability enrichment analysis workflow in 20 LSFs were similar. Each LSF has two types, one is classified within all accessible peaks (84K, aCRE), another input set of peaks are derived from transcribed peaks (19K, tCRE). Firstly, we collected 80 traits to do downstream analysis, only traits with an estimated heritability were carried forward for analysis. (q value >0.2).

For some significant traits, we compared the heritability enrichment level among four conditions (all tCRE, all aCRE, significant group’s tCRE, significant group’s aCRE).

We calculate 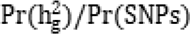 to measure four LSFs’ genetic associations and heritability.

### Jaccard Similarity Analysis

Based on CytoTRACE inference and unique gene expression, the 225 fetal cell types in our study can be grouped into two general categories with respect to cell proliferation. Most differentiated cells, such as cardiac muscle cells in humans, are no longer capable of cell division. These cells are produced during embryonic development, differentiate, and are then retained throughout the life of the organism. In contrast, 54 cell types have been annotated as proliferative state, sustain proliferation.

We obtained each patient’s raw atac counts in those cancer type specific peak sets (https://gdc.cancer.gov/about-data/publications/ATACseq-AWG), filtered out those low detected peaks (reads counts<20) and generate each patient accessible peak set as bed format file, then convert to hg19 chromosome using LiftOver. Meanwhile, we use

bedtools to generate evenly-sized1000-bp bins across genome, score the chromatin accessibility similarity between patients and cell types by calculated Jaccard similarity coefficients using peak signal overlap those windows.

The process to evaluate the cancer type and cell types chromatin state similarity are basically same. We obtain each cancer type specific strongest peak sets and produce a binary bin matrix for cancer and cell types in correspond organ, Jaccard index was computed, and these results were summarized using heatmap.

Then rank these cell types based on the cell type’s Jaccard similarity coefficient, we can evaluate chromatin accessible similarity among 9 malignant cancer types and proliferate cell types by calculate proliferate cell type proportion in top 10% cell types. Specifically, we calculated the hyper-geometric p-value testing the overlap within each cancer’s top10% similar cell types compared to the proliferate cell type set using “phyper” in R. (See TableS7)

---

**Supplementary Fig. 1.**
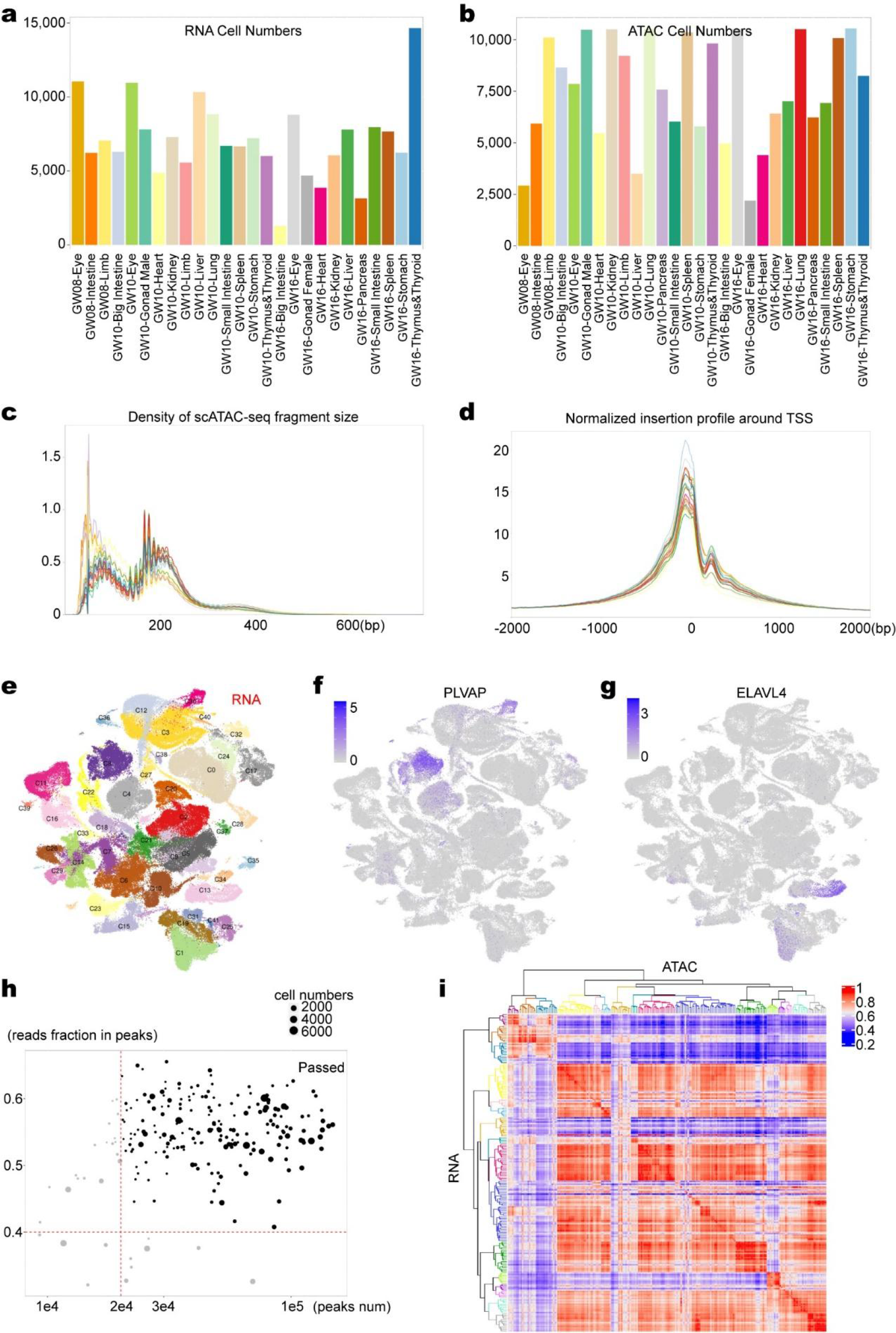
Assessing data quality and validating label transfering result, related to Figure 1. (a,b) Bar charts of cell numbers in each sample in scRNA-seq and scATAC-seq data. (c,d) Left panel: Distribution of sequenced insert sizes for each sample. Right panel: Normalized insertion profile around TSS for each sample. (e-g) (e) UMAP embedding of all 185,061 cells from the scRNA-seq data colored by 41 major clusters. (f) Normalized gene expression level of PLVAP. (g) Normalized gene expression level of ELAVL4. (h) QC of label transferring result. Bubble plot demonstrates the significant peak numbers (x axis), along with read fraction in peaks (y axis) of each cell type in scATAC-seq data. Black dots represent the cell types passing the QC filters. (i) Heatmap of spearman correlations between average gene activity score profiles (x axis) and gene expression profiles (y axis) for 225 cell types. The cell type order is the same as Fig. 1c.

**Supplementary Fig. 2.**
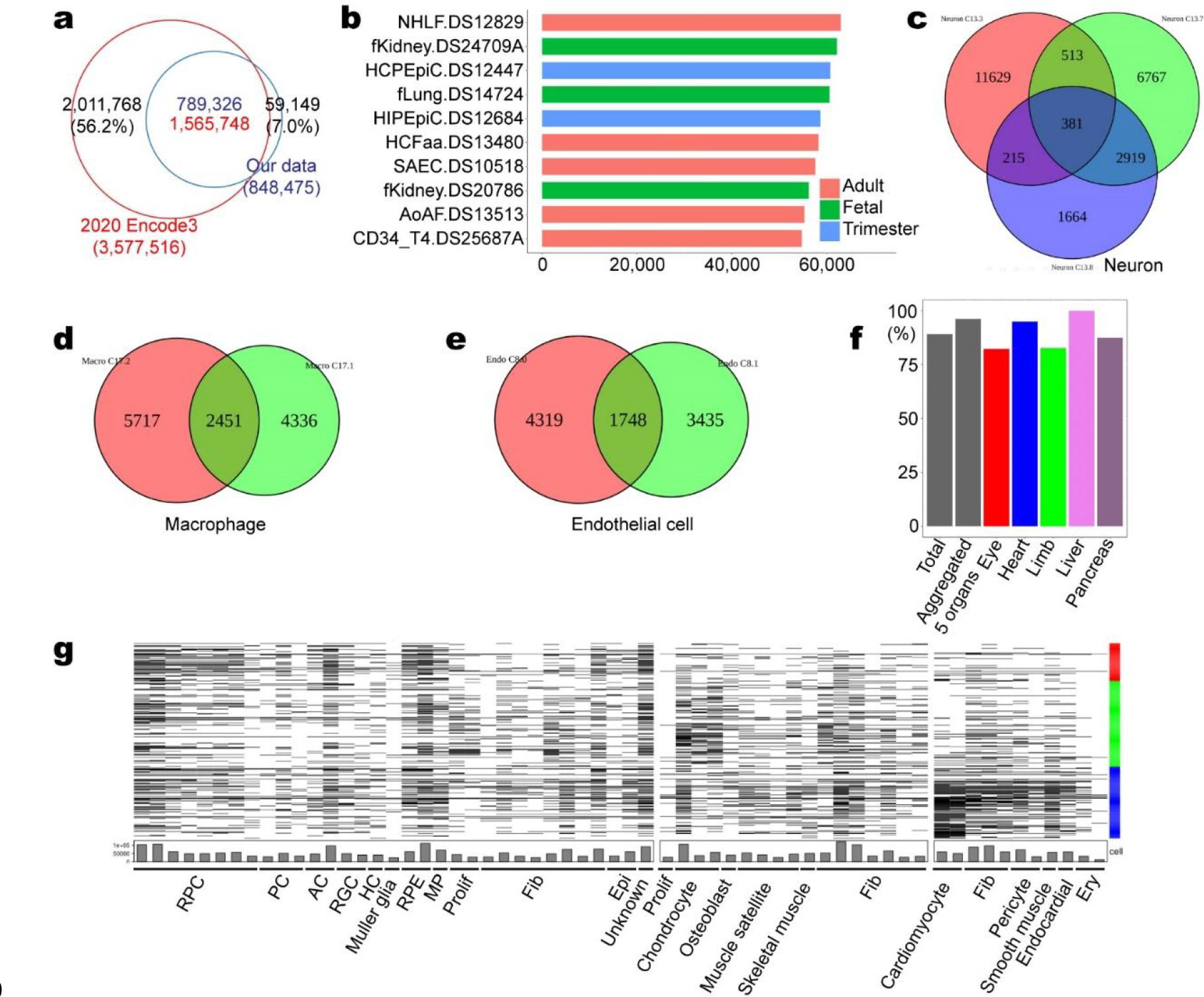
Comparison of chromatin accessible sites, related to Figure 2. (a) The overlap between DHSs from ENCODE3 paper and our ATAC peaks. Using all DHSs. (b) Top 10 tissues that contribute most to DHSs specific peaks (774,300 in Fig. 2b). (c-e) Overlaps of ATAC specific peaks among sub cell types from Fig. 2c. (f) Coverage of VISTA enhancers in different sets. (g) Accessibility of VISTA enhancers among different cell types. Each row represents an enhancer, and each column represents a cell type. The color code on right represents organ source from Supplementary Fig. 2d.

**Supplementary Fig. 3.**
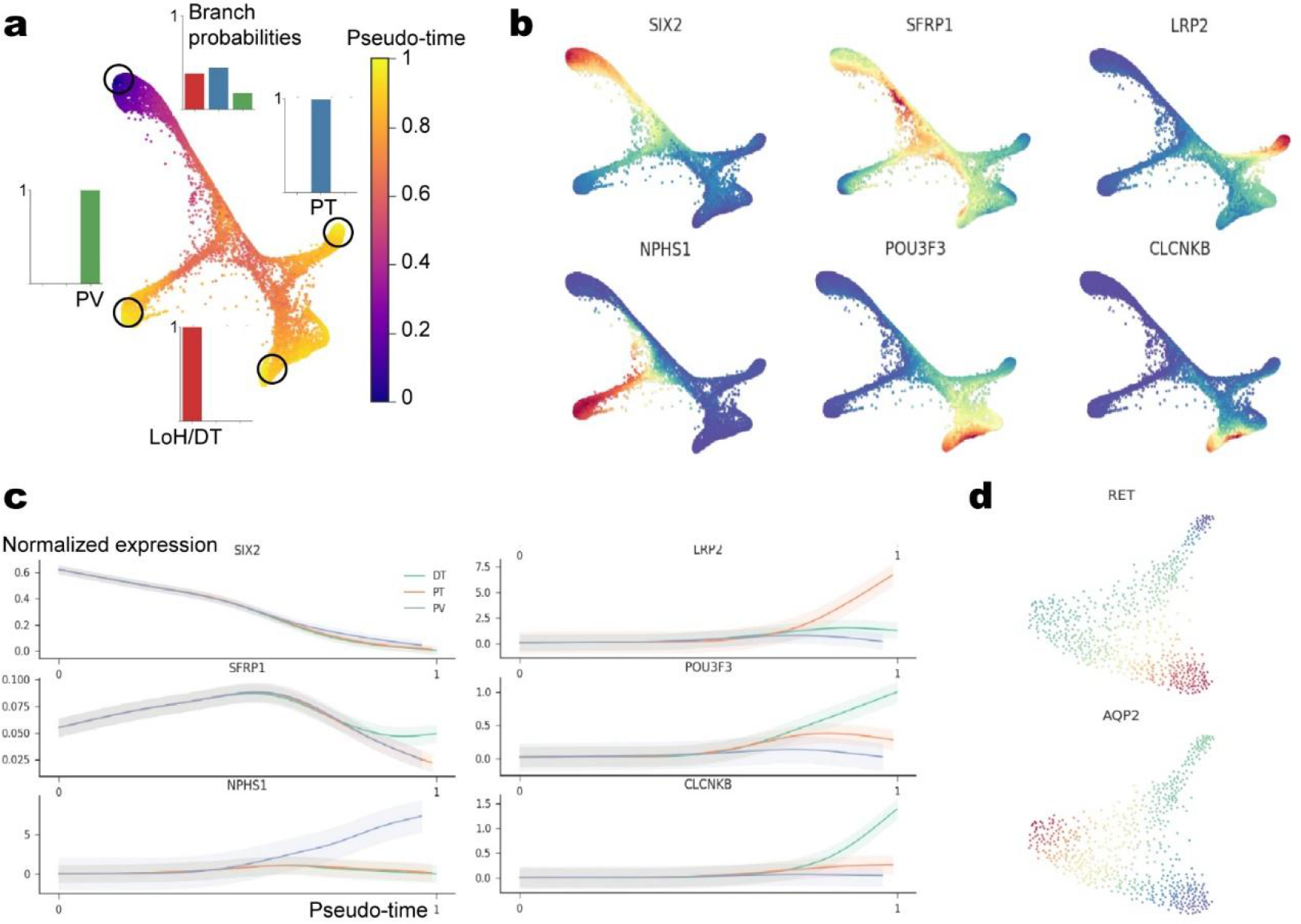
Trajectory analysis of kidney epithelial cells, related to Figure 3. (a) UMAP embedding of CM derived 12,048 cells from the scATAC-seq data colored by pseudo-time. The bar chart shows the terminal state probability distributions of three selected cells. (b) Normalized gene expression level of previous known markers. (c) Expression pattern of previous known marker genes in each segment along the pseudo-time path. (d) UMAP embedding of UB derived 604 cells from the scATAC-seq data colored by expression of UB markers (top) and CD marker (bottom).

**Supplementary Fig. 4.**
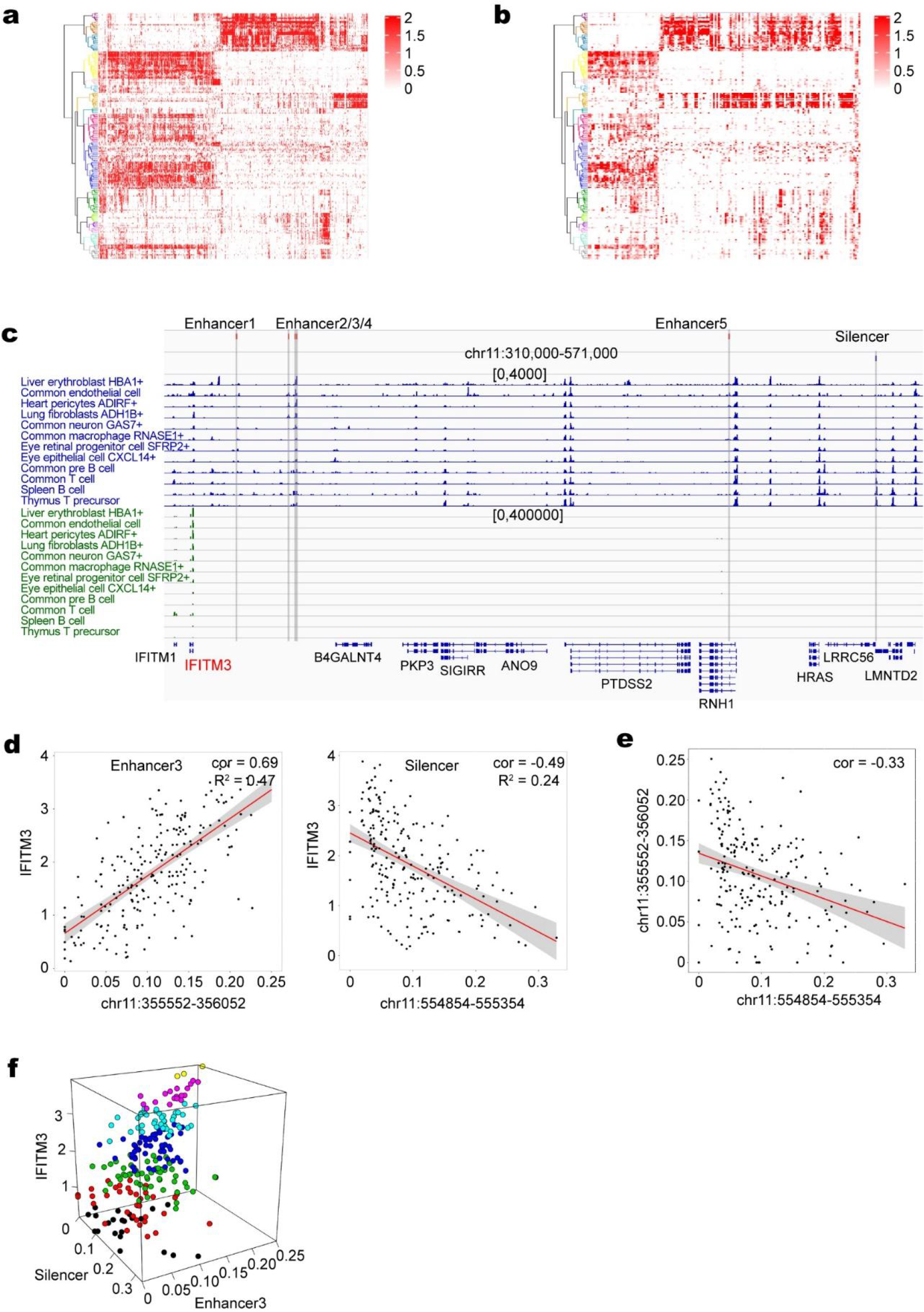
Accessibility of positive and negative cis-regulatory elements, related to Figure 5. (a-b) Heatmaps showing the combinational regulation on the same gene. Same as Fig. 5b,c, but using all enhancers or silencers. (c) Example locus around IFITM3 with annotated cis-regulatory elements on the top. Cell types are ordered according to expression level of IFITM3. (d) Scatter plot demonstrates the peak accessibility level (x axis), along with IFITM3 expression level (y axis) of each cell type. Left is enhancer 3 and right is silencer from (c). (e) Scatter plot demonstrates the accessibility level of the silencer (x axis), along with the accessibility level of the enhancer 3 (y axis) of each cell type. (f) 3D scatter plot showing the relationship among the accessibility level of the enhancer 3, silencer and gene expression of IFITM3.

**Supplementary Fig. 5.**
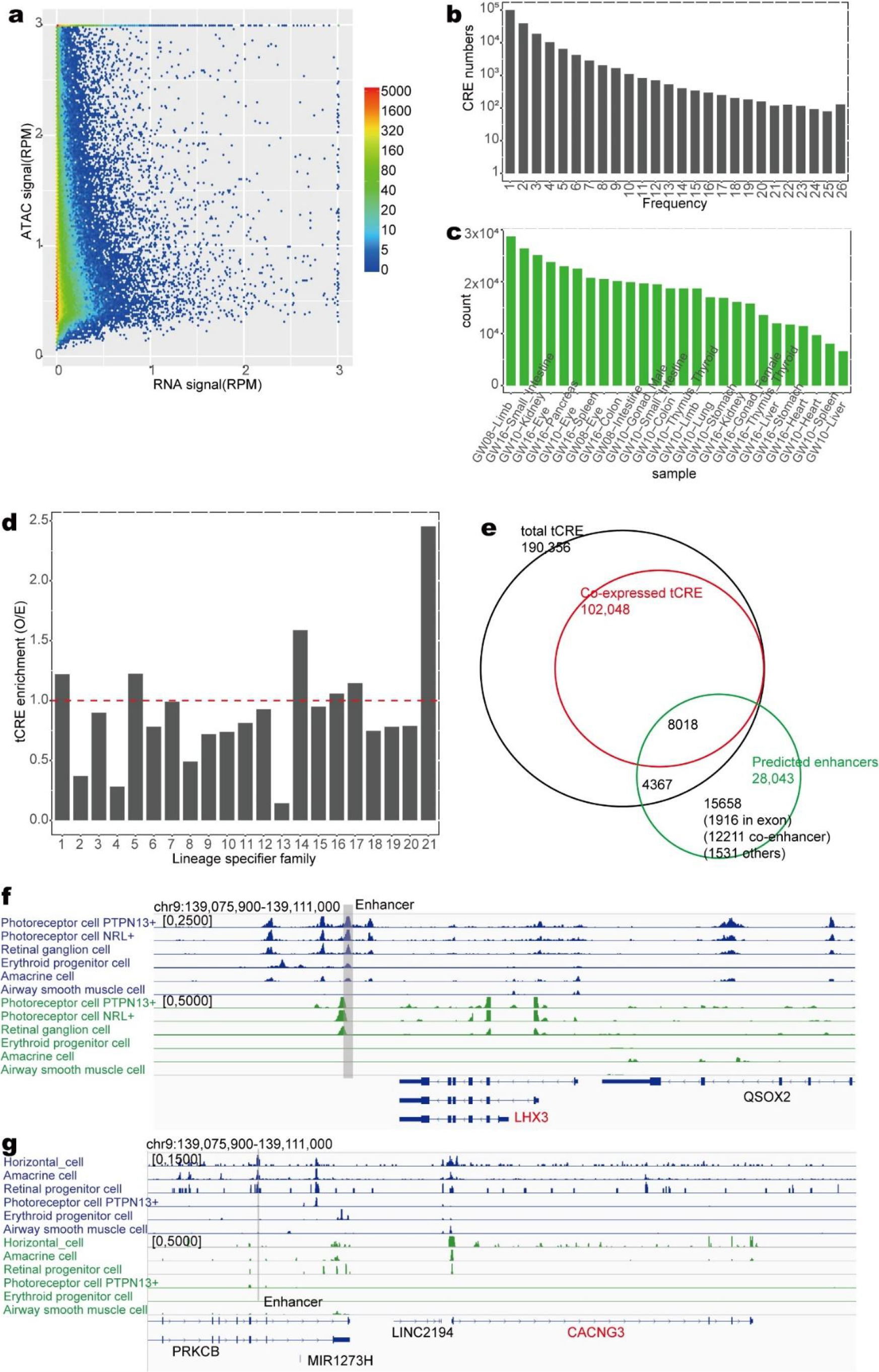
Assessing properties of transcribed cis elements, related to Figure 6. (a) Smooth scatter plot demonstrates the peak transcription level (x axis), along with ATAC signal intensity (y axis) of transcribed cis elements in GW10 limb. Only cis elements with open chromatin state are shown. (b) Frequency distribution of transcribed cis elements in all samples. (c) Counts of identified transcribed cis elements in each sample. (d) Enrichment for transcribed cis elements in different peak groups from Fig 2A. (e) The overlap between transcribed cis elements, co-expressed cis elements and putative enhancers from peak-to-gene links. (f) Example locus of transcription-dependent enhancer of LHX3 with annotated cis-regulatory elements on the top. Cell types are ordered according to expression level of LHX3. (g) Example locus of transcription-independent enhancer of CACNG3 with annotated cis-regulatory elements on the top. Cell types are ordered according to expression level of CACNG3.

**Supplementary Fig. 6.**
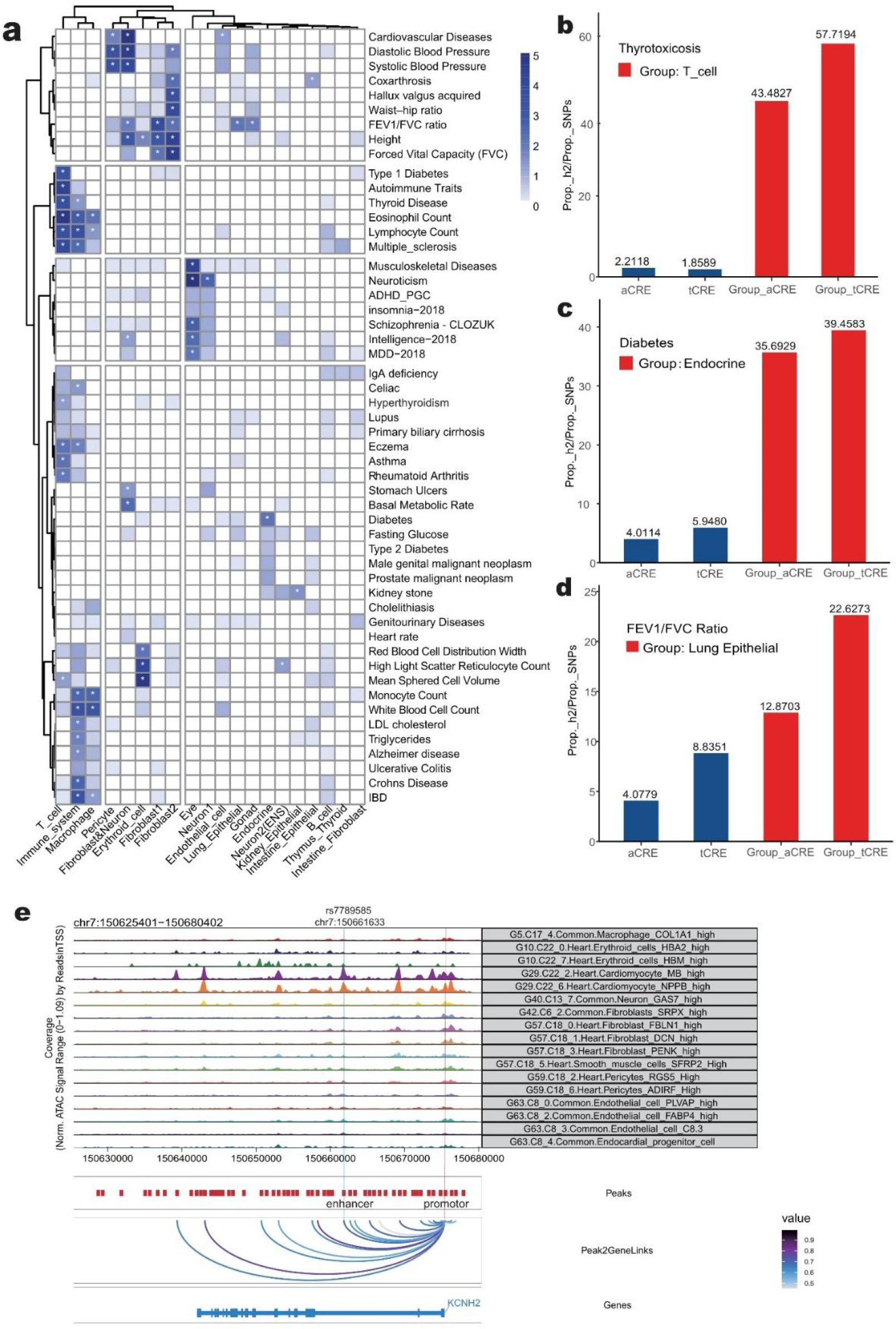
S-LDSC results from 52 traits show heritability enrichment in 20 LSFs. (a) Heatmap displaying the -log10(q value of the enrichment) for 20 peak groups across 52 traits analyzed (Except LSF 21). 20 LSFs were classified and colored by broader cell-type category, that met the across 20 LSFs, significance level (q<0.05) are indicated with an asterisk. (b-d) Bart plots displaying Enrichment of heritability in various CRE-types. X-axis, from left to right are ’aCRE’ represent total cis-elements detected (840K); ’tCRE’ represent total transcribed cis-elements (190K); ’Group_aCRE’ represent specific peak group of total cis-elements; Group_tCRE’ represent specific peak group of transcribed cis-elements. Y-axis, Heritability enrichment Pr(h2)/Pr(SNPs), estimated by LDSC. Red bar shows heritability enrichment of assigned group peak, the blue bar shows bulk level. (e) Genome browser tracks for scATAC-seq (top; scale, RPM) and indicated one AF-associated risk variant. Co-accessibility track shows linkages between the AF variant–containing CRE and promoters.

**Supplementary Fig. 7.**
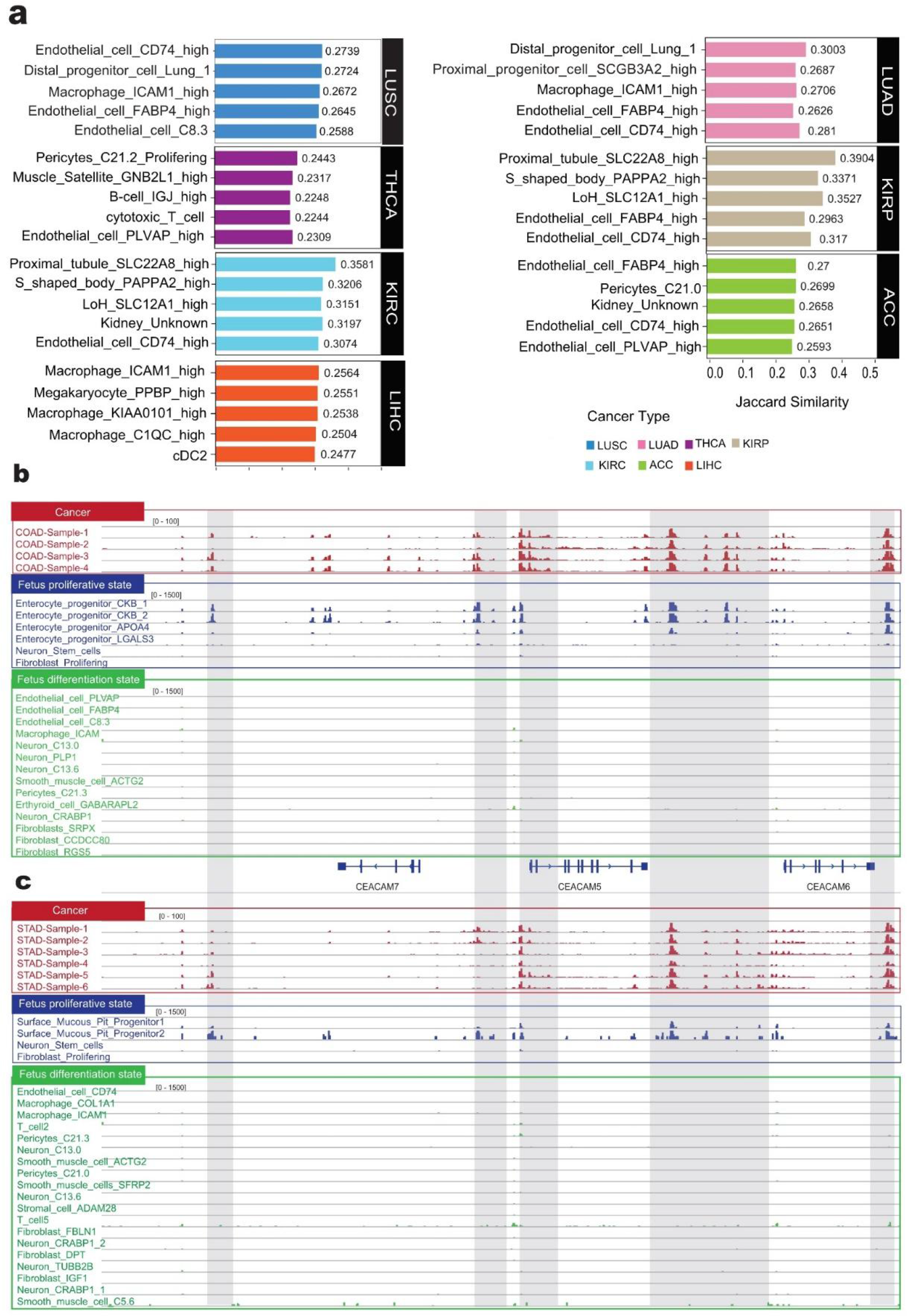
(a) Bar plot showing the Jaccard similarity Score of top5 similar fetal cell types for 7 cancer types (See Supplement table). (b) Regulatory landscape around the CEACAM family genes (CEACAM5, CEACAM6, CEACAM7), indicating GENCODE gene annotations, ATAC seq tracks for each cell type of Colon (blue and green), and top 6 from COAD sample (Red). Fetal cell types in colon have been classified two parts (See Method), those cell types which are labeled by green color belong to differentiation state cell types, other blue cell types are proliferative state cell types. (c) The Same as FigureS7B, highlight the chromatin profile between COAD and Surface Mucous Pit Progenitor cells.

